# A comprehensive examination of Nanopore native RNA sequencing for characterization of complex transcriptomes

**DOI:** 10.1101/574525

**Authors:** Charlotte Soneson, Yao Yao, Anna Bratus-Neuenschwander, Andrea Patrignani, Mark D. Robinson, Shobbir Hussain

## Abstract

A platform for highly parallel direct sequencing of native RNA strands was recently described by Oxford Nanopore Technologies (ONT); in order to assess overall performance in transcript-level investigations, the technology was applied for sequencing sets of synthetic transcripts as well as a yeast transcriptome. However, despite initial efforts it remains crucial to further investigate characteristics of ONT native RNA sequencing when applied to much more complex transcriptomes. Here we thus undertook extensive native RNA sequencing of polyA+ RNA from two human cell lines, and thereby analysed ~5.2 million aligned native RNA reads which consisted of a total of ~4.6 billion bases. To enable informative comparisons, we also performed relevant ONT direct cDNA- and Illumina-sequencing. We find that while native RNA sequencing does enable some of the anticipated advantages, key unexpected aspects hamper its performance, most notably the quite frequent inability to obtain full-length transcripts from single reads, as well as difficulties to unambiguously infer their true transcript of origin. While characterising issues that need to be addressed when investigating more complex transcriptomes, our study highlights that with some defined improvements, native RNA sequencing could be an important addition to the mammalian transcriptomics toolbox.

## Introduction

The extent and observed complexity of cellular mRNA splicing patterns appear to have generally expanded during the course of evolution ^1^, and in more advanced species, several subtly different mRNA transcript isoforms are likely to exist for most genes ^2–4^. Within a biological organism, the observed pattern of mRNA splicing for a given gene also frequently varies between tissues and cell types, and can even respond to external cues or changes to the environment ^5^. Thus, the ability to readily perform transcript-level functional investigations will almost certainly enrich our understanding of a number of important biological processes. To enable this to be accomplished in a reliable manner, methods that can unequivocally distinguish and quantify the presence of transcript isoforms from the raw sequence reads are required.

Recently, long-read sequencing methodologies have been introduced into the transcriptomics field, offering the opportunity to directly generate individual reads that can span the full length of transcripts ^6–12^. This could, for example, ameliorate problems associated with earlier technologies’ needs for DNA-mediated amplification and computational transcript assembly from short sequence reads ^13,14^. Notably, the newer long-read Oxford Nanopore Technologies (ONT) platform now also provides the ability to sequence native RNA strands directly ^15^. In their study, ONT described the efficient use of native RNA sequencing to yield reliable abundance estimates of full-length transcripts from a yeast polyA+ transcriptome as well as sets of standardized synthetic transcripts. However, larger transcriptome sizes, and in particular the much higher complexity of splicing patterns that can be observed in higher organisms, might potentially pose additional challenges during such transcript-level investigations.

With the aim to characterize the gene- and transcript-level composition of complex transcriptomes, in this study, we applied ONT long-read native RNA-sequencing to samples from two human cell lines; HAP1 and HEK293. We also performed matched ONT direct (PCR-free) cDNA sequencing as well as regular Illumina RNA-seq to enable relevant comparisons and assessments. For computational analysis, considering the lower accuracy of Nanopore sequencing, we primarily employed a reference-based approach, estimating abundances of a set of annotated transcript isoforms and genes. An additional motivation for this was that also in situations where a reference-free approach is used for transcript *identification*, reference-based methods are often useful for subsequent *quantification* of transcript abundances. We present our findings relating to differences between the performance of a variety of analysis algorithms, and the potential advantages that current ONT direct RNA-seq brings over the traditional Illumina sequencing, as well as current limitations of the technology.

## Results

### Overall data characteristics

We utilized three distinct ONT library preparation workflows in this study, all having in common that RNA or cDNA molecules are sequenced directly without PCR. For our initial efforts, during which direct cDNA sequencing kits were not available from ONT, we modified the regular ONT-NSK007 2D PCR-based workflow in order to enable 1D direct cDNA sequencing (see *Methods)* (Fig. 1A). We also made use of the subsequently released ONT-DCS108 kit for direct cDNA sequencing, incorporating enrichment for full-length cDNAs (Fig. 1B). Most of the data presented in this study, however, was obtained using the ONT-RNA001 kit for native RNA sequencing (Fig. 1C). All ONT sequencing was performed using R9.4 flow cells, and the ONT Albacore package was used for basecalling. We noted concerns of previous studies reporting that filtering of reads during basecalling often results in a significant number of useful good-quality reads being discarded ^10^. Indeed, in subsequent versions of available ONT Albacore basecalling packages, filtering was either turned off as default or offered as an option. As sequencing depth would likely be the key limiting factor influencing our downstream analyses, and reasoning that true low-quality reads would be filtered during the alignment step, we thus made use of the Albacore non-filtering option.

**Figure 1.**
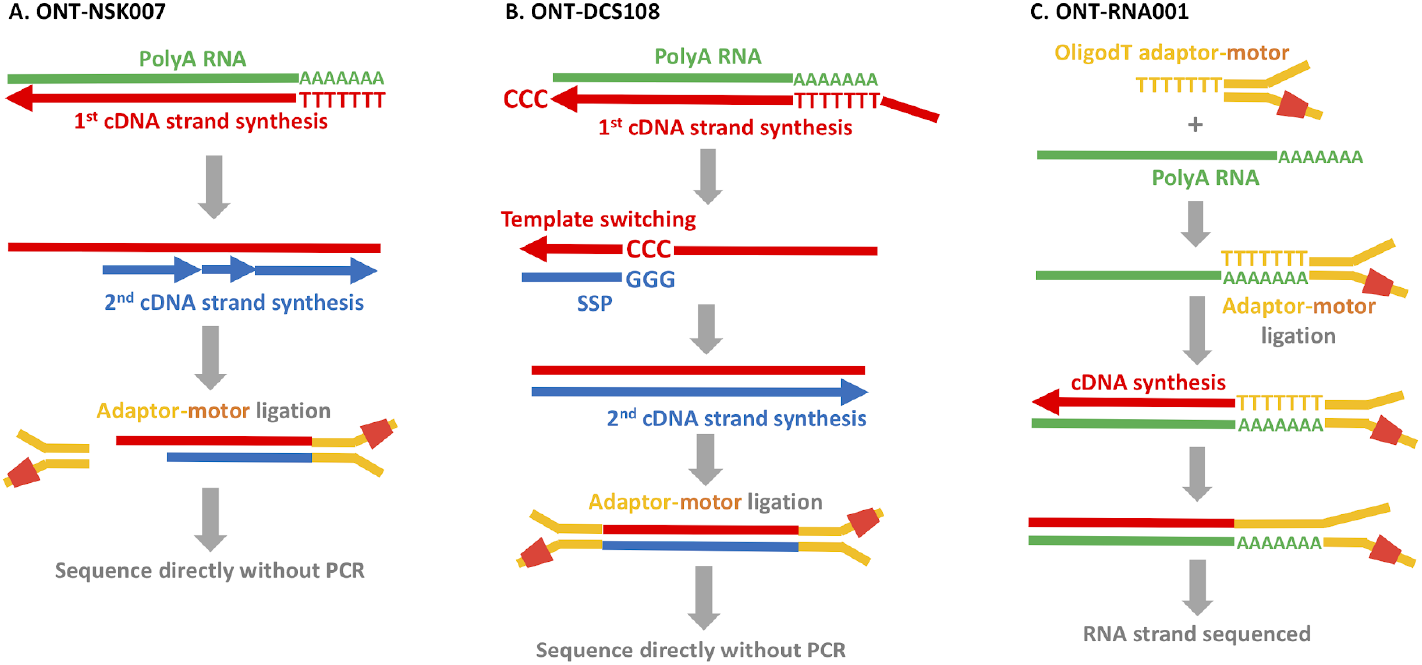
Overview of library preparation workflows used in this study. **A.** In the ONT-NSK007 cDNA library preparation method, polyA RNA is used as a template for first strand cDNA synthesis which is initiated from an oligodT primer. The NEB second strand cDNA synthesis module (E6111) is then used to generate double-stranded cDNAs; here random primers are used to initiate cDNA synthesis, the products of which are stitched together by DNA ligase. Note that since priming of second strand synthesis occurs randomly, as depicted here this may not always begin from the very end of the first strand template. Adaptor-motor complexes are then ligated to the double-stranded cDNA ends prior to direct sequencing (the motor is an enzyme which will feed the nucleic strand into the nanopore). Note that instances where the first strand overhang might be particularly long, as in the example depicted here, it is probably unlikely that the adaptor-motor complex will ligate efficiently to enable sequencing of the second strand, though the first strand will still be sequenced. **B.** To better enrich for full length cDNAs, the ONT-DCS108 direct cDNA sequencing kit, which leverages the template switching phenomenon ^16^, was used. When the first strand cDNA synthesis reaches the end on the RNA molecule, the reverse transcriptase will add a few non-template Cs to the end of the cDNA. A Strand Switching Primer (SSP) present in the reaction binds to these non-templated Cs, and the reverse transcriptase then switches template from the RNA to the SSP. The second cDNA strand, presuming its synthesis continues to the end of the first strand template, will also span the full length of the primary polyA RNA template. Following adaptor ligation, the double stranded cDNAs are then sequenced directly. **C.** The ONT-RNA001 workflow enables sequencing of native RNA strands. Here an oligodT-adaptor-motor complex is ligated to the polyA end of the RNA. In order to relax the secondary structure of the RNA (and thus help ensure efficient translocation of the RNA strand through the nanopore), a cDNA synthesis step is performed. Since only the RNA strand has a motor ligated, the RNA molecule but not the cDNA strand is always sequenced.

The yield from the different ONT protocols varied between approximately 500,000 and 1,500,000 unfiltered reads per sample (Supplementary Fig. 1A), and the read length distributions were overall similar among the libraries, with a peak close to 1,000 bases (Supplementary Fig. 1B). The distribution of average base qualities per read varied between the different types of libraries (Supplementary Fig. 1C), with cDNA libraries as expected ^8^ showing higher base qualities than native RNA libraries. We also noticed an association between the read length and the average base quality, with both very short and very long reads often having lower quality (Supplementary Fig. 2).

### Genome and transcriptome alignment

The ONT reads were aligned to the human reference genome and transcriptome using minimap2 (see *Methods).* The N50 values for the portion of a read aligned to the genome were 907, 1,210, 1,043 and 941 bases for the *ONT-NSK007-HAP, ONT-DCS108-HAP, ONT-RNA001-HAP*, and *ONT-RNA001-HEK* data sets, respectively (median aligned lengths for the respective data sets were 633, 765, 621 and 596 bases, and the longest aligned read parts were 75,756, 20,681, 12,839 and 14,692 bases in the four data sets). As we aligned unfiltered reads, the alignment rates across library types were unsurprisingly only modest, varying between 55 and 76% for the genome alignment, and from 45 to 73% for the transcriptome alignment (Fig. 2A). As expected, the unaligned reads were enriched for low base qualities (Supplementary Fig. 3A), and thus largely represented reads that would have been classified as ‘failed’ during automatic filtering. In comparison, for the four matching Illumina libraries, STAR aligned between 89 and 94% of the reads uniquely to the genome, with an additional 2-2.5% of the reads aligning in multiple locations. The *ONT-DCS108-HAP* libraries showed the largest differences between the genome and transcriptome alignment rates (60-69% vs 45-51%), whereas the rates for the other data sets were more similar. It is possible that one explanation for the large difference in genomic and transcriptomic alignment rates for this data set could be a contamination with genomic DNA. In the *ONT-DCS108-HAP* libraries, compared to the set of all reads aligning to the genome, the reads aligning *exclusively* to the genome showed a slight enrichment for long reads with a lower average base quality but where a larger portion of the read aligned. This pattern, however, was not reproduced in the other data sets, where the reads aligning exclusively to the genome were rather shorter and showed a poorer agreement with the reference (Supplementary Fig. 3).

**Figure 2.**
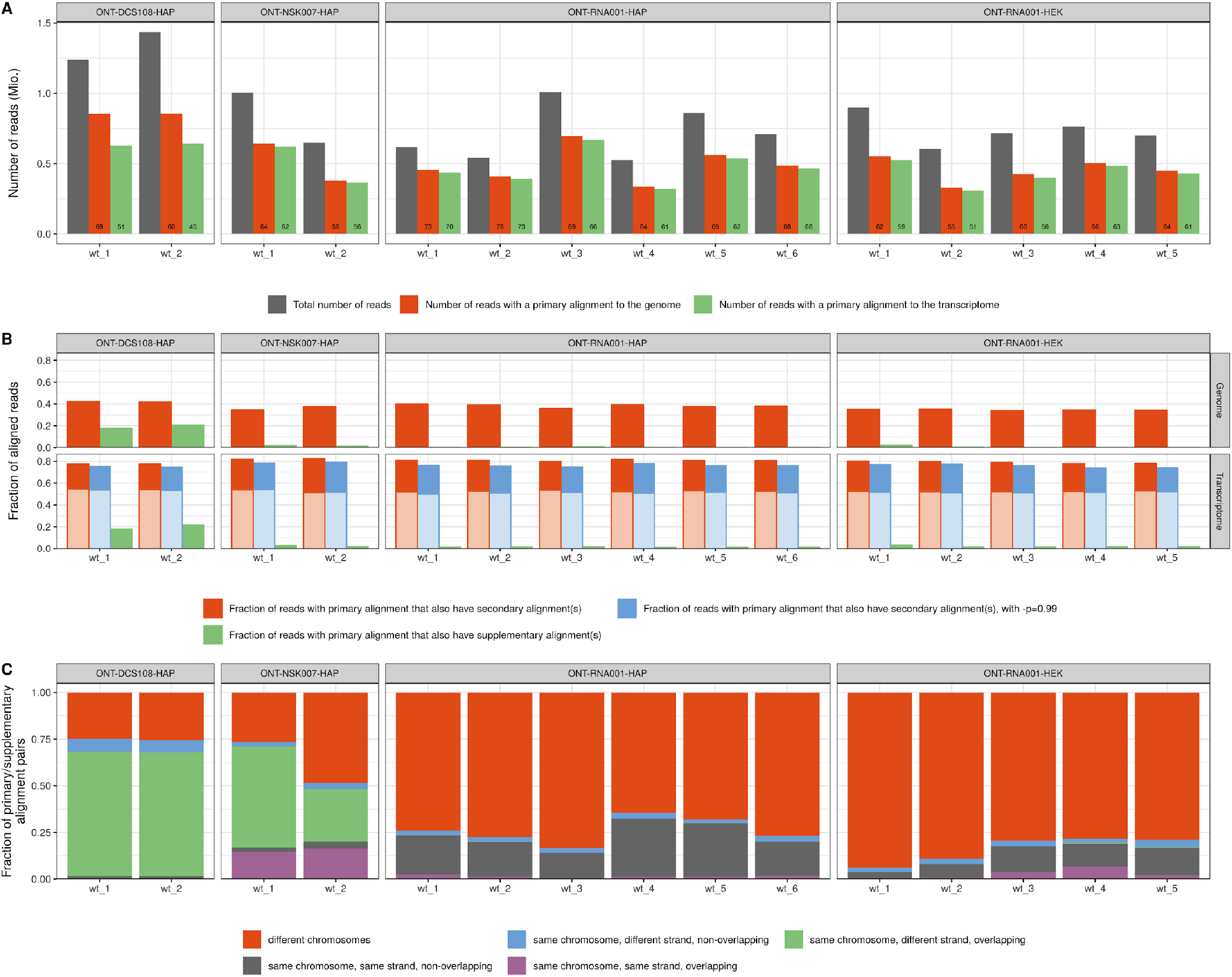
Characterization of aligned reads. **A.** Total number of reads and the number of reads with a primary alignment to the genome or transcriptome, respectively, in each of the ONT libraries. The number displayed in each bar represents the alignment rate in % (the fraction of the total number of reads for which minimap2 reports a primary alignment). **B.** Fraction of the reads with a primary alignment to the genome or transcriptome, respectively, that also have at least one reported secondary or supplementary alignment. The lighter shaded parts of the secondary transcriptome alignment bars correspond to reads where all primary and secondary alignments are to isoforms of the same gene, while the darker shaded parts correspond to reads with reported alignments to transcripts from different genes. **C.** Investigation of supplementary genome alignments. Each supplementary alignment is categorized based on whether it is on the same chromosome and strand as the primary alignment, and if the alignment positions of the primary and supplementary alignments overlap.

Approximately 40% of the reads with a primary genome alignment could be mapped to multiple places in the genome, i.e., had also at least one reported secondary genome alignment (Fig. 2B). For most libraries, a single secondary alignment was most common, while for the *ONT-DCS108-HAP* libraries, a larger fraction of reads had more than five secondary genome alignments (Supplementary Fig. 4A). As expected, due to the high similarity among transcripts, the fraction of reads with at least one secondary alignment increased to approximately 80% for the transcriptome alignment (Fig. 2B). Again, a small number of secondary alignments was most common (Supplementary Fig. 4B). The secondary alignment rate was only marginally affected by increasing the -p argument of minimap2, which sets the minimal accepted ratio between the alignment score of secondary and primary alignments, to 0.99 instead of the default 0.8 (Fig. 2B). For a majority of the reads, the target transcripts of all primary and secondary transcriptome alignments were isoforms of the same gene (Fig. 2B), suggesting that the main source of ambiguity is on the individual isoform level rather than on the gene level. Only a small part of the secondary alignments (typically less than 5% of the reads) arose due to the presence of multiple fully identical transcripts in the Ensembl reference catalog; in all remaining cases there was at least some difference between the target transcripts of the reported primary and secondary alignments. ‘Unavoidable’ secondary alignments may also be the result of reads stemming from reference transcripts that are proper subsequences of other reference transcripts. Among the 1,044,960 possible pairs of reference transcripts annotated to the same gene in our annotation catalog, there are 64,437 such pairs (6.2%). In these situations, in theory, a read could still be considered ‘unambiguously assignable’ to the shorter transcript if it is similar enough, under the assumption that all ONT reads represent full-length transcripts. Without this strong assumption, effective automated disambiguation would require a reliable model of the read generation process, accounting for the probability of fragmentation of RNA or cDNA molecules in the library preparation step and/or read truncation during the sequencing-basecalling process. To investigate to what extent the secondary alignments in our libraries could be the result of nested sets of reference transcripts, we extracted all reads with at least one secondary transcriptome alignment, and among all primary and secondary alignments, we selected the one for which the covered portion of the target transcript by the read was highest. If the secondary alignments are the result of the true transcript of origin being contained in the other target transcripts, we expect this maximally covered portion to be close to 1. Interestingly, for all the data sets except *ONT-NSK007-HAP*, while there is a clear peak close to 1, there is also a broad distribution of lower coverage degrees (Supplementary Fig. 5). The large number of secondary transcriptome alignments with alignment scores similar to the reported primary alignment suggests that, despite the long read length, unambiguously inferring the true transcript of origin for any given read is still highly non-trivial, and simply selecting the reported primary alignment for downstream analysis can give misleading results.

While secondary alignments represent possible mapping positions of a read beyond the one reported in the primary alignment, *supplementary* alignments arise when a read cannot be mapped in a contiguous fashion, and consequently minimap2 splits the alignment into multiple parts. We observed a comparatively large number of supplementary alignments in the *ONT-DCS108-HAP* data set, both for genome and transcriptome alignments (Fig. 2B). Further investigation revealed that in this data set, as well as in *ONT-NSK007-HAP*, a relatively large fraction of the supplementary alignments overlapped the corresponding primary alignment, but on the opposite strand (Fig. 2C). This observation is interesting, as we note that the ONT ‘1D ^2^’ sequencing mode (https://nanoporetech.com/) exploits the observation that the second strand (which also has a motor enzyme attached) of a double-stranded DNA molecule often enters the sequencing nanopore immediately following the first strand during 1D sequencing. 1D ^2^ sequencing chemistry is designed to further promote this observed phenomenon, and the associated 1D ^2^ base-caller is specifically designed to efficiently split reads according to each strand sequenced. Thus our findings of frequent overlapping supplementary alignments on opposite strands may reflect un-split reads by the standard 1D basecaller. Accordingly, this type of self-chimeric supplementary alignments were almost completely absent in the native RNA samples where single strands, as opposed to double-stranded cDNAs, are present in libraries. For the *ONT-NSK007-HAP* libraries, where often only one of the strands of the double-stranded cDNAs will have a motor enzyme attached (Fig. 1), the relative frequency of this type of supplementary alignments was somewhat lower than in the *ONT-DCS108-HAP* libraries (in addition to the total rate of supplementary alignments begin considerably lower), adding further support to this speculated cause.

A peak of short low-quality unfiltered reads was consistently observed in the native RNA libraries (Supplementary Fig. 1B), and the majority of these did not align adequately to either the genome or the transcriptome (Supplementary Fig. 3A-B). More generally, for aligned reads, in particular those shorter than 10,000 bases, most of the individual bases could be matched to a position in the reference sequence, indicated by a large fraction of “M”s and consequently a low fraction of insertions, deletions and soft-clipped bases in the CIGAR string (Fig. 3, Supplementary Fig. 3C-D). Reads longer than 10,000 bases, which were mostly found in the cDNA libraries, typically did not align end-to-end (Fig. 3). For the *ONT-DCS108-HAP* libraries, a large fraction of the bases in the primary alignments were soft-clipped, corresponding to the large number of supplementary alignments discussed above.

**Figure 3.**
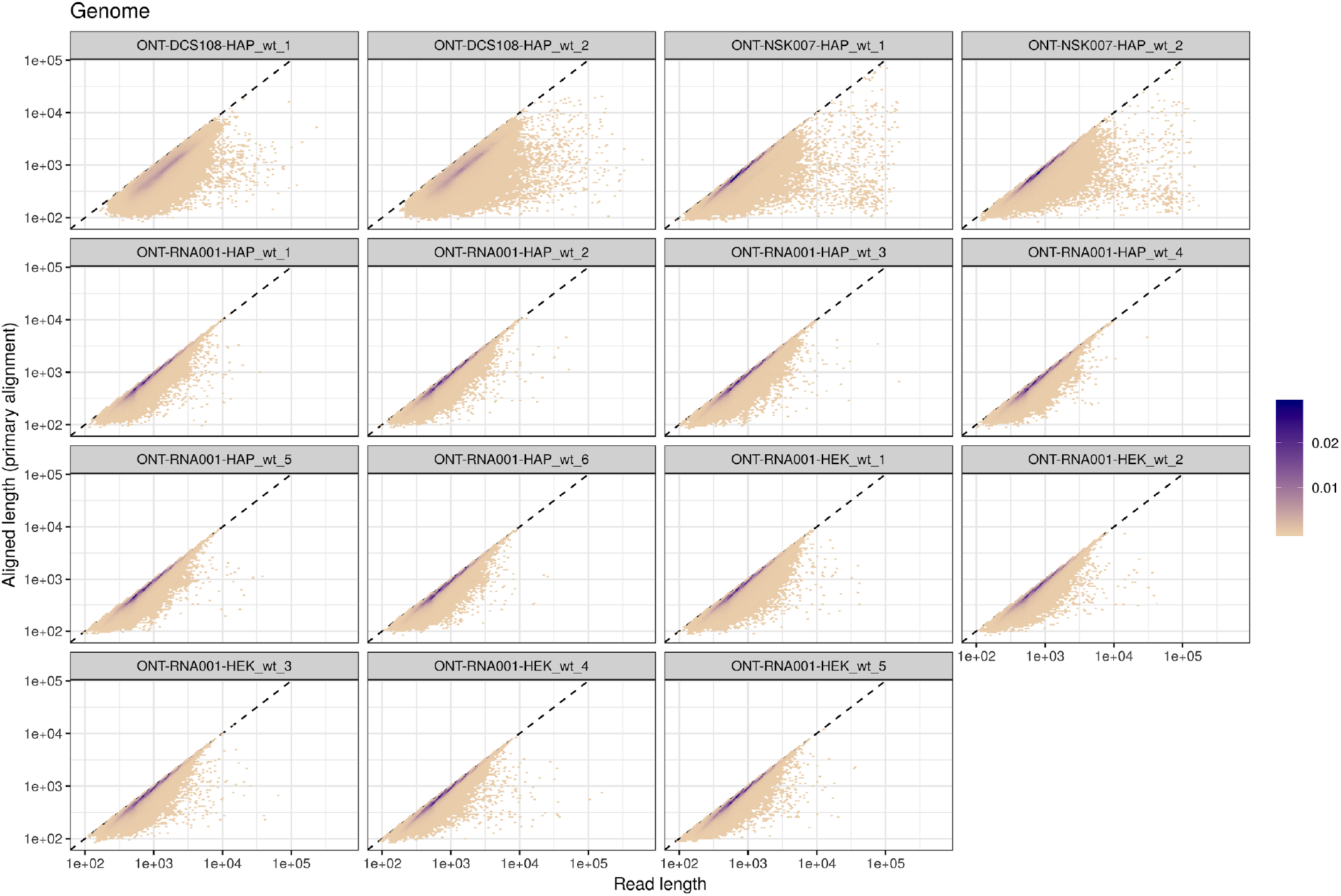
Total read length (*x*) vs aligned length (*y*, the sum of the number of “M” and “I” characters in the CIGAR string) for the primary genome alignment of each read, in each of the ONT libraries. The colour indicates point density.

Incorporating the genomic coordinates of the annotated genes, we observed differences in the gene body read coverage distribution between the libraries (Supplementary Fig. 6), with a stronger 3’ coverage bias in the cDNA libraries than in the native RNA libraries. While given the nature of the library preparation this was expected for the NSK007 cDNA libraries, it is also quite possible that the template switching mechanism does not work to full efficiency in the DCS108 cDNA protocol.

### Coverage of full-length transcripts by individual ONT reads

To investigate to what extent individual ONT reads could be expected to represent full-length transcripts, we selected the “best” target transcript for each read, starting from the set of all primary and secondary transcriptome alignments obtained with minimap2, with -p set to 0.99. For each read, we kept all alignments for which the number of aligned nucleotides was at least 90% of the maximal such number across all alignments for the read, and among these, we selected the one with the largest transcript coverage degree (number of “M” and “D” characters in the CIGAR string of the alignment, divided by the annotated transcript length). While this alignment does not necessarily represent the “true” origin of the read, the procedure gives an upper bound of the degree of transcript coverage achieved by individual reads. As expected from the *ONT-NSK007-HAP* library preparation, which does not involve full-length cDNA enrichment (Fig. 1), reads from this sample achieved a lower degree of full-length transcript coverage across the range of transcript lengths (Fig. 4A). While shorter transcripts could often be completely covered by a single read in the *ONT-RNA001-HAP, ONT-RNA001-HEK* and *ONT-DCS108-HAP* libraries, this was rarely the case for long transcripts (Fig. 4A, Supplementary Figure 7). This observation, that many of the raw ONT reads do not appear to represent full-length transcripts, needs to be taken into account during transcript identification and quantification. Applying the same procedure to the SIRV and ERCC data sets from Garalde *et al.* ^15^ revealed that a majority of these synthetic transcripts were well covered by single reads (Fig. 4B-C), confirming observations from previous studies ^9,15^; importantly, however, all transcripts in the SIRV and ERCC catalogs are shorter than 2,500 bases. In the Ensembl GRCh38.90 catalog, approximately 17% of the transcripts are longer than that, and the coverage degree of these transcripts by single reads were generally less than 50%. This suggests that while the synthetic transcript catalogs provide useful information about the performance of long-read transcriptome sequencing and analysis methods, extrapolation of the results to real, complex transcriptomes should be done with care.

**Figure 4.**
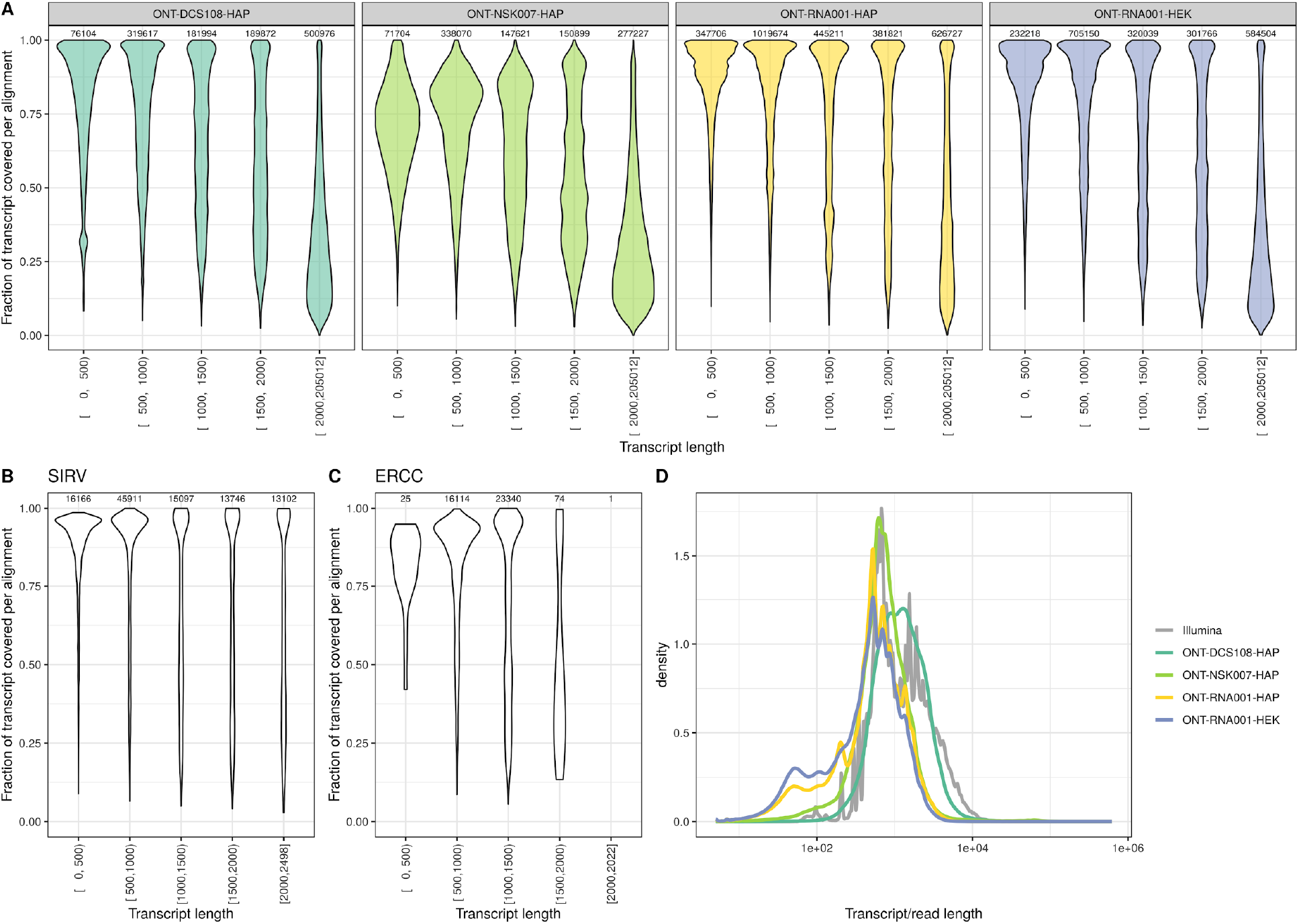
Transcript coverage fraction by individual reads. **A.** Distribution of coverage fractions of transcripts by individual reads, for each of the four ONT data sets, stratified by the length of the target transcript. The ‘target transcript’ was selected to maximize the coverage fraction, among all reported long enough alignments (see text), and thus the reported coverage fractions represent upper bounds of the true ones. The number above each violin indicates the number of processed alignments to transcripts in the corresponding length category. **B-C.** Distribution of coverage fractions of transcripts by individual reads for the SIRV and ERCC data sets. **D.** Observed distribution of raw read lengths (for ONT data sets) and expected distribution of transcript molecule lengths based on annotated transcript lengths and estimated abundances in the Illumina samples. Values are aggregated across all samples within each data set.

To further investigate the degree to which individual ONT reads are likely to represent full-length transcripts, we compared the observed raw ONT read length distribution with the ‘expected’ transcript length distribution in these samples, obtained by weighting the annotated transcript lengths by the estimated transcript abundances (in transcripts per million – TPM) estimated by Salmon in the Illumina samples. This analysis showed an apparent shortage of ONT reads in the length range of the longest transcripts inferred to be expressed in the Illumina data (Fig. 4D). The *ONT-DCS108-HAP* samples were the exception; however, for many of the reads in these libraries, the primary alignment does not cover the entire read (Fig. 3). This holds true both for reads with a supplementary alignment and for those without one (Supplementary Fig. 8A-C). Further inspection of the longer annotated transcripts with high estimated abundances in the Illumina samples revealed consistent base pair coverage by Illumina reads along the length of these transcripts (Supplementary Fig. 8D), indicating that these were indeed likely to be truly present in samples. Moreover, these transcripts were from ‘standard’ genes in that the vast majority of the long transcripts with high estimated abundance in the Illumina samples were annotated as protein coding, and they were found on almost all chromosomes. Overall, such observations further illustrate that using current library preparation and sequencing workflows, long transcripts are often not represented by single ONT reads.

### Reference-based transcript detection and abundance quantification

Four reference-based methods were used to estimate transcript and gene abundances in each of the ONT libraries. For two of these methods, we specifically evaluated the impact of data preprocessing: for minimap2 followed by Salmon in alignment-based mode (denoted salmonminimap2), we investigated the effect of setting the `-p` argument of minimap2 to different values (the default of 0.8 as well as 0.99) in the transcriptome alignment step, and for Salmon in quasi-mapping mode, we evaluated the effect of providing only the aligned bases of the reads with a primary alignment anywhere in the genome (see *Methods).* Increasing -p to 0.99 led to a slightly improved correlation between ONT transcript read counts and estimated transcript abundances from the Illumina samples (obtained by Salmon in quasi-mapping mode), and thus, in the following analyses, we set -p equal to 0.99 for Salmon following minimap2 (Supplementary Fig. 9). Removing the non-aligned bases before running Salmon did not improve the correlations notably (Supplementary Fig. 9). Since this is a more involved procedure, and further introduces a dependency on the genome alignments, we use the Salmon quantifications obtained using the original, non-truncated reads for the rest of the analyses.

We observed a large difference between the numbers of reads that were assigned to features by the different quantification methods (Supplementary Fig. 10). The highest assignment rates were consistently obtained with salmonminimap2, where all reads that were aligned to the transcriptome were also subsequently assigned to features. featureCounts assigned a slightly lower fraction of the reads to genes, while Salmon in quasi-mapping mode and Wub assigned considerably fewer reads. However, the relatively low number of reads assigned by Salmon in quasi-mapping mode were distributed across as many, sometimes more, genes and transcripts as the reads assigned by salmonminimap2 (Fig. 5A-B), suggesting that no category of genes or transcripts was consistently missed. In general, the transcript-level detection rate increased with transcript length, both for ONT and Illumina libraries (Fig. 5C). Counting the number of “detected” transcripts and genes, defined as the number of features with an expected read count of at least 1 with salmonminimap2 (ONT) or Salmon (Illumina), at various degrees of subsampling (Fig. 5D-E) suggested that the current sequencing depth of approximately 0.5 million mapped ONT reads per library was not enough to detect all expressed genes or transcripts. Furthermore, the number of observed genes were similar to the number observed in the Illumina libraries if these were subsampled to comparable sequencing depths. With the aim of investigating whether there are systematic ‘blind spots’ in the detection of features in the ONT data (in which case we expect the same set of transcripts to be detected in all libraries) or if the lack of saturation is purely a result of undersampling (in which case we would expect differences in the set of detected transcripts across libraries), we compared the saturation curves obtained from individual samples to that obtained by first pooling the reads across all replicates within a data set, and subsequently sampling from this pool (Supplementary Fig. 11). On the transcript level, pooling the samples improved the degree of saturation for a given number of reads, while no improvement could be seen on the gene level.

**Figure 5.**
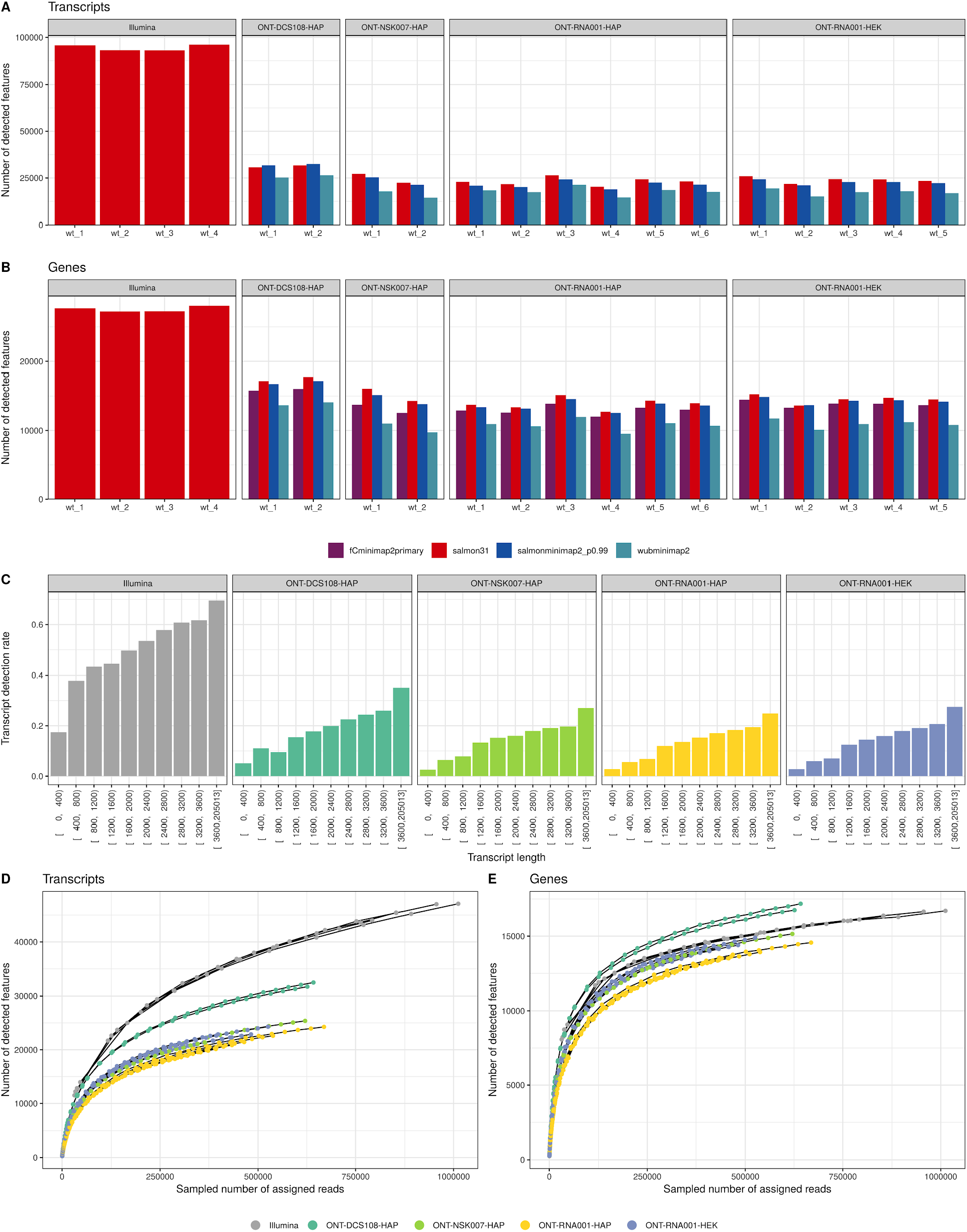
Detection of annotated transcripts and genes. **A-B**. Number of detected transcripts and genes with the applied abundance estimation methods, in each library. Here, a feature is considered detected if the estimated read count is ≥1. **C**. Fraction of transcripts detected (with estimated count ≥1) in at least one sample, stratified by transcript length, in the respective data sets. **D-E**. Saturation of transcript and gene detection, in ONT and Illumina libraries. For each library, we subsampled the reads and recorded the number of transcripts and genes detected with an estimated salmonminimap2 count (ONT libraries) or Salmon count (Illumina libraries) ≥1. The Illumina curves are truncated to the range of read numbers observed in the ONT libraries.

Next, we calculated the correlation between abundance estimates among replicates of the HAP cell line, within and between data sets. As expected, the correlation between replicates was higher on the gene level than on the transcript level, and higher within a data set than between data sets (Supplementary Fig. 12). On the gene level, correlation between replicates was almost as high in the ONT data as in the Illumina data, for all quantification methods, while for transcript-level abundances, higher correlations were observed in the Illumina data. Overall, Wub showed the highest correlation of abundance estimates between replicates in the ONT data sets. Notably, correlations between cDNA and native RNA samples were as high as those among samples obtained with different cDNA protocols.

Comparing the abundance estimates obtained for the same library with different quantification methods showed that, perhaps unsurprisingly, Salmon in quasi-mapping mode and salmonminimap2 had the highest correlation (Supplementary Fig. 13). Stratifying transcripts and genes by the annotated biotype suggested that certain biotypes (in particular, short transcripts such as miRNAs) were consistently assigned very low abundances with ONT, while they were observed in the Illumina libraries (Supplementary Fig. 14).

### Transcript identifiability

Next, we focused on specific transcriptomic features that are useful for discriminating similar isoforms. First, we extracted the junctions observed after aligning the ONT reads to the genome. The majority of the junctions that were covered by at least 5 ONT reads were already annotated in the reference transcriptome, while this was more rarely the case for lowly-covered junctions (Fig. 6A-B). Junctions that were observed in the ONT reads but did not correspond to annotated junctions were less likely than those already annotated to be observed in the Illumina data, and also less likely to harbor a canonical splice junction motif (GT-AG) (Supplementary Fig. 15). Not surprisingly, individual ONT reads generally spanned more junctions than Illumina reads (Supplementary Fig. 16), which should provide improved ability of correct transcript identification.

**Figure 6.**
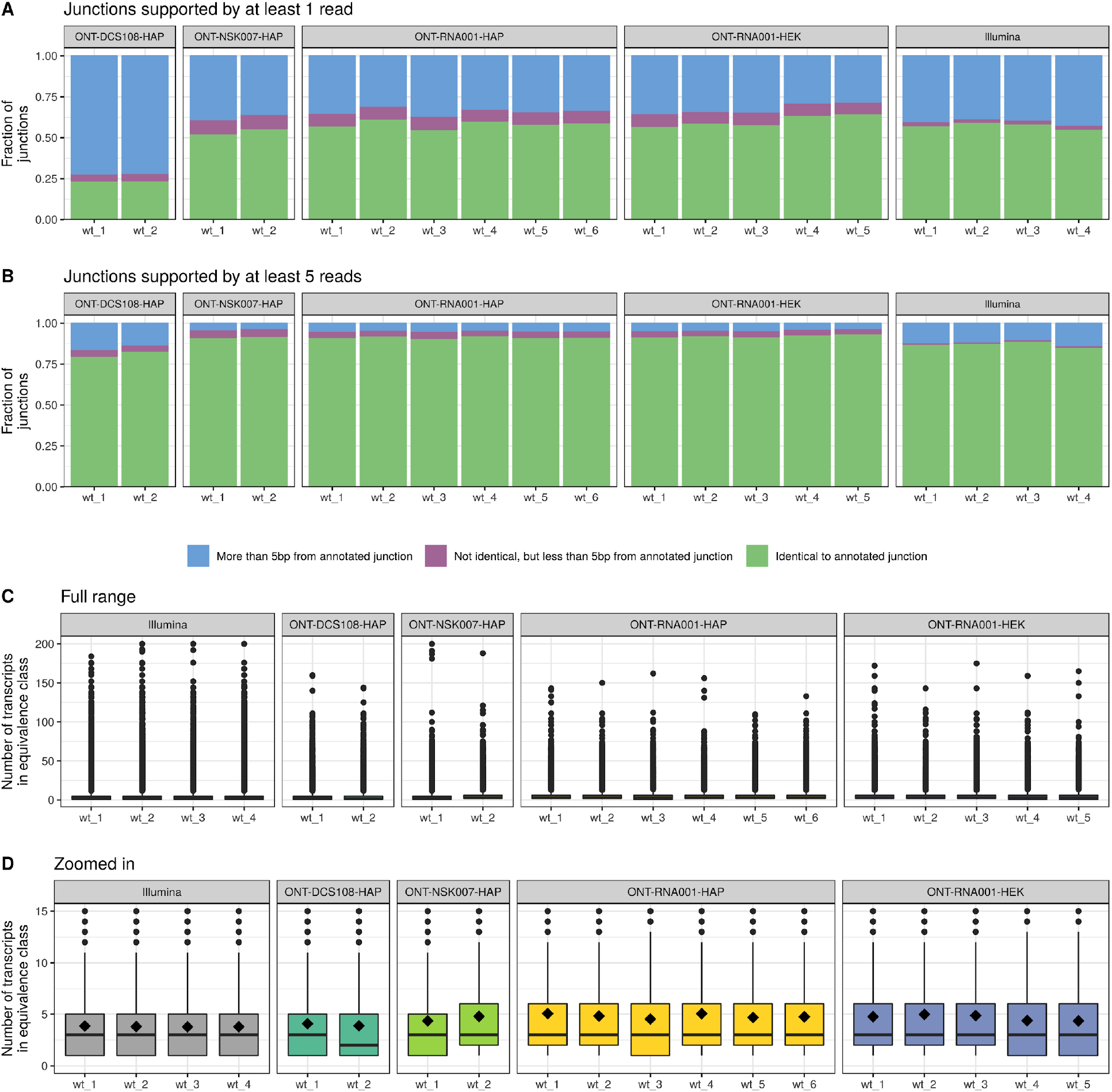
**A-B**. Annotation status of junctions observed in each ONT and Illumina library. A junction is considered observed if it is supported by at least 1 **(A)** or 5 **(B)** reads. For each observed junction, the distance to each annotated junction was defined as the absolute difference between the start positions plus the absolute difference between the end positions. This distance was used to find the closest annotated junction. **C**. Distribution of the number of transcripts contained in the Salmon equivalence class that a read is assigned to, across all reads, for each ONT and Illumina library. **D**. As **C**, but zoomed in to the range [0, 15]. The black diamond shape indicates the mean.

In order to further investigate if the longer length of ONT reads compared to Illumina reads in fact improved their unambiguous assignment to specific transcripts, we tabulated the number of transcripts included in the equivalence class that each read was assigned to when running Salmon in quasi-mapping mode. A read being assigned to a large equivalence class indicates that the read sequence is compatible with many annotated transcripts, and consequently that unambiguous assignment is difficult. While fewer ONT reads were assigned to equivalence classes with a very large number of transcripts compared to the Illumina counterparts, the average number of transcripts in the equivalence class, across all reads, was almost identical for the ONT and Illumina libraries (Fig. 6C-D). To investigate to what extent this was an effect of the high redundancy among the annotated transcripts, we ran Salmon with the same index, but using the annotated transcript catalog as a proxy for error-free, full-length ‘reads’. In this case, 87% of the reads were assigned to equivalence classes containing a single transcript. This illustrates both that even in this idealized situation, not all reads would be unambiguously assignable to a single annotated transcript, due to redundancies in the annotation catalog, and that for our ONT reads, the ambiguity is still considerably higher than in the ideal situation. Together with the large number of secondary transcriptome alignments observed above, this illustrates the challenging nature of reference-based transcript identification based on ONT reads. Furthermore, the read generation model used by Salmon is adapted to Illumina reads, and thus is likely suboptimal for inferring transcript abundances from ONT reads.

### Reference-free transcript identification

In addition to the reference-based transcript identification and quantification discussed above, we generated a set of high-confidence consensus transcripts for each ONT data set using FLAIR (https://github.com/BrooksLabUCSC/flair). For this analysis, only reads with a 5’ end close to a known promoter region were considered, and only transcript sequences supported by at least 3 ONT reads were retained. The identified transcripts from FLAIR were compared to the annotated reference transcriptome using gffcompare (https://ccb.jhu.edu/software/stringtie/gffcompare.shtml). This comparison identified the most similar reference transcript for each FLAIR transcript that showed at least some overlap with the reference transcriptome, and further assigned a class code describing the type of relationship to this most similar reference transcript (see https://ccb.jhu.edu/software/stringtie/gffcompare.shtml for a description of all class codes). Interestingly, only a relatively low fraction of the identified transcripts in each data set contained a junction chain that was identical to that of an annotated transcript (Fig. 7A, class code ‘=’), while a larger fraction of the identified transcripts contained a junction chain that was consistent with an annotated transcript, but only contained a subset of the junctions. This corroborates the previous observations that many ONT reads may not represent full-length transcript sequences. There is a marked difference compared to the set of transcripts assembled with StringTie from the Illumina samples, a larger fraction of which contain a complete intron chain match with an annotated transcript. There is also a larger fraction of Illumina-derived transcripts that do not overlap known transcripts (Fig. 7A, class code ‘u’). FLAIR transcripts with a junction chain perfectly matching an annotated transcript (class code ‘=’) spanned a range of lengths and number of junctions (Fig. 7B-C), suggesting that transcript identification is not limited to, e.g., short isoforms. Overall, the set of transcripts assembled by StringTie from the Illumina data were more often multi-exonic than those from the ONT libraries, and also spanned a broader range of transcript lengths. A random selection of FLAIR transcript sequences (from the *ONT-RNA001-HAP* library) corresponding to annotated transcripts are shown in Supplementary Fig. 17, to illustrate the variety of transcripts that could be identified.

**Figure 7.**
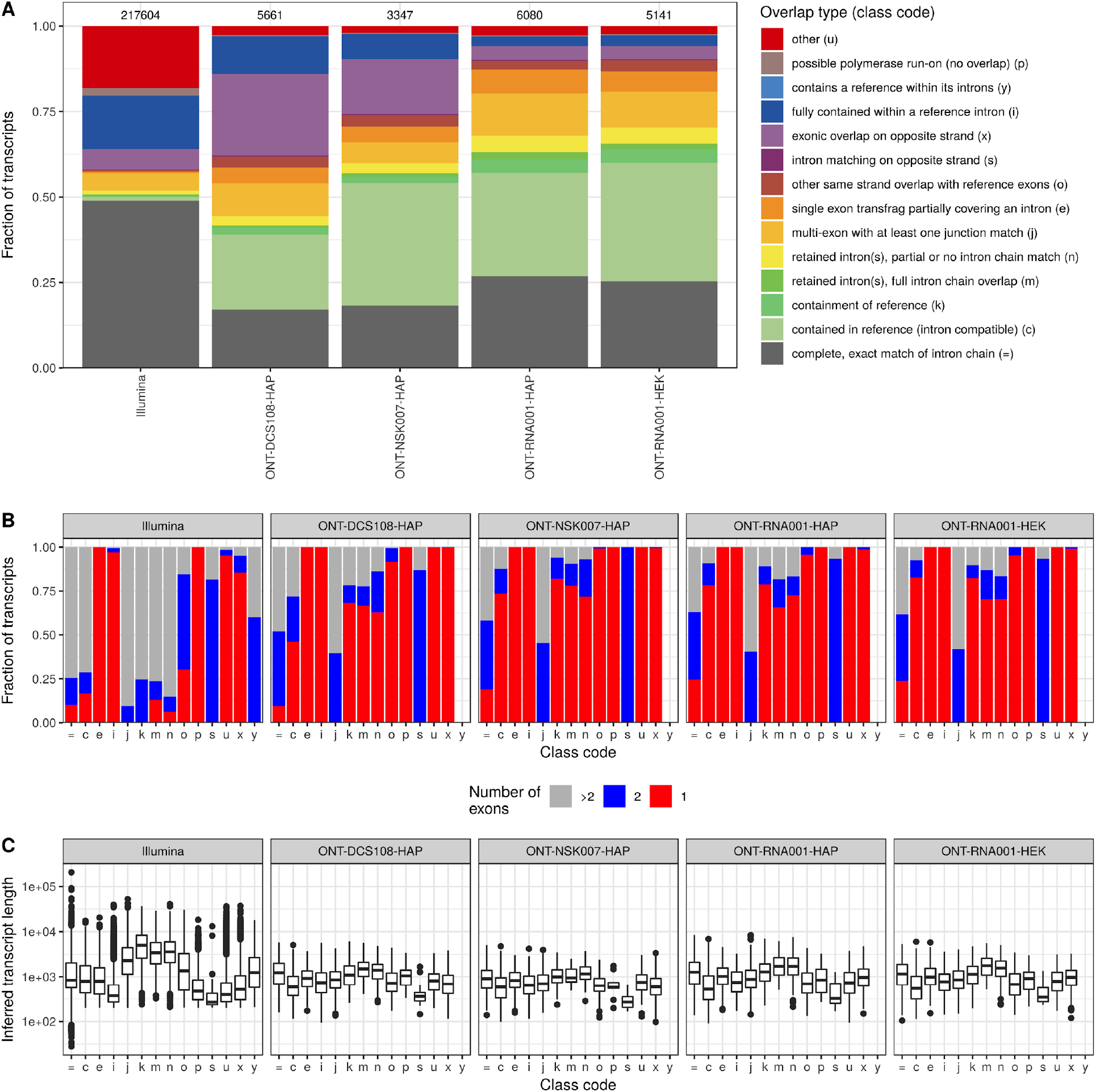
Characterization of transcripts identified by FLAIR. **A**. Class code distribution for de novo identified transcripts from FLAIR (for ONT libraries) or StringTie (for Illumina libraries), compared to the set of annotated transcripts using gffcompare. The number above each bar represents the number of assembled transcripts. The class code for a transcript indicates its relation to the closest annotated transcript. **B**. Number of exons in each transcript identified by FLAIR/StringTie, stratified by the relation to the annotated transcripts (represented by the assigned class code). **C**. Length distribution of transcripts identified by FLAIR/StringTie, stratified by the relation to the annotated transcripts (represented by the assigned class code).

Comparing the set of annotated reference transcripts that could be identified by at least one FLAIR transcript (class code ‘=’ or ‘c’) in the respective ONT data sets showed that a large fraction of these transcripts were only identified in a single data set (Fig. 8A). In addition, reference transcripts identified by the native RNA-sequencing protocol in the two different cell lines showed a higher degree of similarity to each other than to those identified with the cDNA protocols in the HAP cell line, suggesting that transcript identification can be strongly affected by the library preparation protocol. Of note, the native RNA protocols provide information about the strandedness of the reads, which is not the case for the cDNA protocols employed here. Reference transcripts with junction chains corresponding to at least one FLAIR transcript generally showed a higher expression level in the Illumina samples than the reference transcripts that were not identified in any ONT data set (Fig. 8B), suggesting that one possible explanation for the discrepancy between the transcripts identified in the different ONT data sets could be the limited sequencing depth, and that a larger number of ONT reads may be necessary to identify a stable set of expressed transcripts.

**Figure 8.**
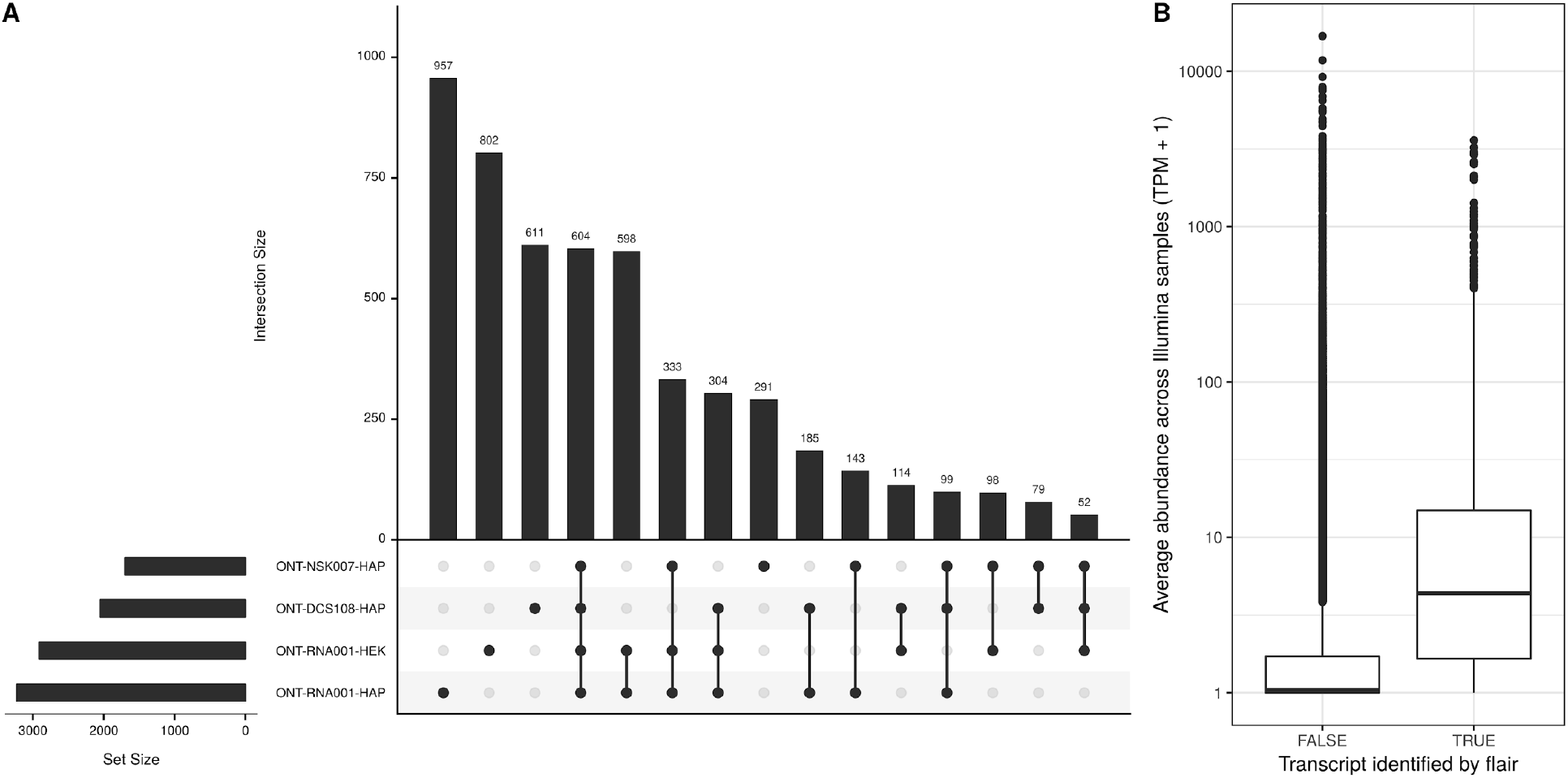
Comparison of annotated transcripts identified by FLAIR in the four ONT data sets. **A.** UpSet plot representing overlaps between the annotated transcripts that are identified by FLAIR in the different ONT data sets. An annotated transcript is considered to be identified if at least one FLAIR transcript is assigned to it with a class code of either ‘=’ or ‘c’. These sets of annotated transcripts are then compared between data sets. Horizontal bars indicate the total number of identified annotated transcripts in the respective data sets, and vertical bars represent the size of each intersection of one or more sets of identified transcripts. **B.** Average abundance across the Illumina samples, for annotated transcripts that are considered ‘identified’ or not by FLAIR. An annotated transcript is considered to be identified if at least one FLAIR transcript from at least one data set is assigned to it with a class code of either ‘=’ or ‘c’.

## Discussion

We have performed a detailed evaluation of reads from Nanopore native RNA sequencing as well as complementary direct cDNA sequencing, from the perspective of transcript identification and quantification. The libraries were prepared from human cell lines, which adds a level of complexity compared to many previous studies focusing on either less complex model organisms or synthetic transcripts. In addition, matched Illumina data was generated for comparison.

We observed that despite the fact that ONT reads are around an order of magnitude longer than typical Illumina reads, identification of their transcript of origin is still highly nontrivial, and a large number of secondary transcriptome alignments with mapping scores very close to the primary alignments were observed for all libraries. This suggests that quantification methods that focus exclusively on the reported primary alignment are likely to be suboptimal, and can be highly biased depending on how the primary alignment is selected among a set of equally-good mappings. We expect that reference-based transcript abundance estimation methods that are able to incorporate information about these multi-mapping reads are more likely to produce reliable abundance estimates; however, to our knowledge no ONT-specific such method, with a read generation model adapted to the ONT library generation, currently exists.

*De novo* as well as reference-based identification of transcripts suggested that a considerable number of the raw ONT reads are likely to not represent full-length reference transcripts. This can have implications for transcript identification and quantification. For example, it is difficult to determine whether a truly truncated version of a reference transcript is present in a sample, or if the reads rather are fragments of a longer transcript molecule. In addition, by attempting to mitigate this issue, e.g. by filtering the ONT reads to only retain those that overlap a known promoter region, the quantitative nature of the data, as well as the number of usable reads, may be reduced. While our manuscript was in preparation for submission, a preprint authored by Workman and colleagues ^17^ was published that highlighted some of key benefits of Nanopore native RNA sequencing, but also indeed reported the very frequent presence of truncated Nanopore reads in native RNA libraries. They were also able to estimate that a significant proportion of transcripts may be truncated by nanopore signal noise, caused for example by electrical signals associated with motor enzyme stalls or by otherwise stray current spikes of unknown origin. These surprising findings are supported by our observations that single native RNA reads frequently fail to cover the full length of transcripts. We also agree that nanopore native RNA read truncation is unlikely due to some fundamental limitation of nanopore-based sequencing, especially considering that ONT 1D genomic DNA sequence reads of several kilobases are consistently achieved without issue using the current pore-type ^18–20^ used to sequence both DNA and RNA. Further, such problems could conceivably be addressed, to at least some extent, by training basecallers to reliably recognize relevant nanopore signal noise events which might cause single molecule sequence reads to be truncated or split.

An inability to read approximately 10-15 nucleotides at the 5’ end of each strand, and relatively higher error rates, were identified as the two principal drawbacks of Nanopore native RNA sequencing by the Workman et al study, although these are potentially readily addressable ^17^. Here we highlight that the sequencing depths achieved from native RNA libraries, typically ~0.5M aligned reads per flow cell, are likely not enough to saturate transcript detection, either using reference-based or *de novo* approaches. Further, our attempts at relevant differential expression analyses from native RNA sequencing data during a parallel study suffered from low power and high variability (data not shown), most likely due to the limited coverage within each library replicate. Improving throughput (the amount of sequence rendered per unit cost and unit time) is a critical issue: if ONT sequencing throughput remains low, uptake and thus impact within the transcriptomics field will likely remain limited, even given its distinguished benefits. Although protein-pore sequencing can be scaled to considerably higher levels (i.e. either on the ONT GridION or PromethION instruments), the associated consumable nanopore array costs remain high. Thus, native RNA-seq throughput characteristics that are deemed acceptable by the transcriptomics community at large will likely require a highly-optimized RNA motor enzyme, or ultimately a shift to a lower cost nanopore array type. When characterization of complex transcriptomes at transcript-level comprises the project remit, our study here describes that Nanopore direct RNA-seq remains a roundly promising but fledgling analysis tool.

## Methods

### Cell lines and culture

HEK293 cells (ATCC) were cultured in Dulbecco’s Modified Eagles Medium (DMEM) supplemented with 10% FBS and penicillin/streptomycin. HAP1 cells (Horizon Discovery) were grown in Iscove’s Modified Dulbecco’s Medium (IMDM) supplemented with 10% FBS and penicillin/streptomycin. All cultures were maintained at a temperature of 37°C in a humidified incubator with 5% CO2. When required, exponentially growing cells were harvested by washing in Phosphate Buffered Saline (PBS) and then incubating with Trypsin-EDTA, followed by further washing of pelleted cells in PBS.

### Library preparation and sequencing

For the Nanopore libraries, total RNA was extracted from cell pellets using Trizol, and the polyA+ fraction isolated using oligodT dynabeads (Invitrogen). The ONT kits NSK007, DCS108, and RNA001 were then used for PCR-free 1D library preparations. For RNA001, 500ng of input polyA+ RNA was used per sample and the libraries were made following ONT instructions. For DCS108, 100ng of input polyA+ RNA was used per sample and the libraries were prepared according to ONT instructions. For NSK007, 100 ng of input polyA+ RNA was used per sample and libraries were made according to ONT instructions, except that the hairpin adaptor (HPA) ligation and PCR steps were omitted as described previously ^18^, in order to enable 1D and direct cDNA sequencing respectively. The prepared libraries were sequenced on the MinION using R9.4 flow cells with the relevant MinKNOW script to generate fast5 files. All generated fast5 reads were then basecalled in Albacore (version 1.2 for NSK007 libraries and version 2.1 for DCS108 and RNA001 libraries) using the relevant script to yield fastq files. As Albacore only contained a 2D script for NSK007 basecalling, only the generated NSK007 fastq ‘raw’ reads (i.e. complement and template) were taken forward for analysis, while any attempted ‘consensus’ reads present were discarded.

For the Illumina samples, all libraries were made using the Illumina TruSeq stranded mRNA kit. The mRNA libraries were prepared from 500 ng of Trizol-extracted total RNA using the Illumina TruSeq^®^ Stranded mRNA Sample Preparation Kit with 15 PCR cycles applied. Libraries were quantified and quality checked using qPCR with Illumina adapter specific primers and Agilent 2200 TapeStation, respectively. Diluted indexed mRNA-seq (10nM) libraries were pooled, used for cluster generation (Illumina TruSeq PE Cluster Kit v4-cBot-HS) and sequenced [Illumina HiSeq 4000, Illumina TruSeq SBS Kit v4-HS reagents, paired-end approach (2×150bp) with 40-55 million reads per sample].

### Genome and transcriptome alignment

ONT reads were aligned to the human genome (Ensembl primary assembly GRCh38) and transcriptome (combined cDNA and ncRNA reference fasta files from Ensembl GRCh38.90) using minimap2 v2.12 ^21^. The genome alignments were performed with the arguments `-ax splice -N 10`, to allow spliced alignments and up to 10 secondary alignments per read. Alignment files from minimap2 were converted to bam format, sorted and indexed using samtools v1.6 ^22^. The Bioconductor package GenomicAlignments (v1.32.0) ^23^ was used to extract junctions from the alignments. For each observed junction, we calculated the distance (the absolute difference between the start positions plus the absolute difference between the end positions) to the closest annotated junction. For the transcriptome alignment, we used the arguments `-ax map-ont -N 100` to allow more secondary alignments, given the high similarity among transcript isoforms. The minimap2 -p argument, representing the minimal ratio of the secondary to primary alignment score that is allowed in order to report the secondary mapping, has a default value of 0.8. For transcriptome alignment, we investigated the effect of increasing this value in order to restrict the number of reported “suboptimal” secondary alignments. To evaluate the alignments, we recorded the alignment rates, defined as the fraction of reads with a reported primary alignment, as well as the aligned fraction of each read, which we defined as the sum of the number of “M”and “I” characters in the CIGAR string, divided by the full length of the read. For some reads, minimap2 also reported supplementary alignments. For each supplementary genome alignment, we compared the alignment position to that of the corresponding primary alignment, and recorded whether these were on the same or different chromosome and/or strand, and whether the primary and supplementary alignments overlapped each other. Finally, we generated reduced FASTQ files by retaining only reads with a primary alignment to the genome, and for each such read, we removed all bases that were (soft-)clipped in the primary alignment. The resulting bam files were converted to FASTQ format using bedtools bamtofastq v2.27.0 ^24^, and the reads were subsequently shuffled using bbmap v38.02 (https://sourceforge.net/projects/bbmap/). RSeQC v2.6.5 ^25^ was used to examine the coverage profile along gene bodies for each library, based on the GENCODE basic v24 bed file downloaded from https://sourceforge.net/projects/rseqc/files/BED/Human_Homo_sapiens/ on October 23, 2018.

### Gene and transcript abundance estimation

Four different computational methods were used to estimate transcript and gene abundances for the ONT libraries. First, we applied Salmon v0.11.0 ^26^ in quasi-mapping mode, with an index generated from the combined Ensembl cDNA and ncRNA reference fasta files and using the default k value of 31 (denoted salmon31 below). For comparability across pipelines, we retained any duplicate transcripts in the index generation. The mean and maximal fragment lengths were set to 600 and 230,000, respectively, and the flag `—dumpEq` was set to retain equivalence class information. Salmon was also run in quasi-mapping mode on the modified FASTQ files, containing only the aligned part of the primary alignments as described above. Second, we applied Salmon in alignment-based mode to the output bam files from the minimap2 transcriptome alignment, using the flag `—noErrorModei` to disable the default short-read error model of Salmon in the quantification (denoted salmonminimap2). Third, we applied the bam_count_reads.py script from the Wub package (https://github.com/nanoporetech/wub) to the output files from the transcriptome alignment, setting the minimal mapping quality (-a argument) to 5 (denoted wubminimap2). Finally, we applied featureCounts (from subread v1.6.0) ^27,28^ to the primary genome alignments, requiring a minimum overlap of 10 bases and using the `-L` argument to enable the long-read mode (denoted fCminimap2primary). While the Salmon variants and Wub provided transcript-level abundance estimates, which were also aggregated to the gene level, featureCounts provided only gene-level counts and was therefore not considered for transcript quantification.

### De novo transcript identification

In addition to the reference-based quantification described above, we also performed reference-free, *de novo* transcript identification using FLAIR (obtained from https://github.com/BrooksLabUCSC/flair on December 16, 2018), applied to the combined primary genome alignments from all libraries in each ONT data set. The minimap2 bam files were converted to bed format using the bam2bed12.py script provided with FLAIR, and identified junctions were subsequently corrected by comparison to the reference annotation, using the default window size of 10. Next, the corrected reads were collapsed using FLAIR, requiring that the 5’ end of the read falls close to a promoter and retaining only transcripts represented by at least 3 reads. The promoter bed file was obtained by combining active, weak and poised promoters identified in nine cell lines by the ENCODE consortium (obtained from https://genome.ucsc.edu/cgi-bin/hgFileUi?db=hg19&g=wgEncodeBroadHmm and lifted over to hg38 coordinates using the UCSC Genome Browser liftOver tool) The identified transcripts from each data set were compared to the annotated transcripts using gffcompare (https://ccb.jhu.edu/software/stringtie/gffcompare.shtml), whereby each FLAIR transcript was assigned a *class code*, detailing the way in which it is related to the most similar reference transcript.

### Processing of Illumina libraries

Sequencing adapters were removed from the Illumina libraries with TrimGalore! v0.4.4 (http://www.bioinformatics.babraham.ac.uk/projects/trim_galore/, using cutadapt v1.13 ^29^), with quality and length cutoffs both set to 20, and aligned to the Ensembl GRCh38.90 primary genome assembly using STAR v2.5.1b ^30^. Abundances of annotated transcripts were estimated using two different methods: first, with StringTie v1.3.3b ^31^ using reads aligned with HISAT2 v2.1.0 ^32^ (with the `--dta` flag set and using a known splice site file), and second, with Salmon in quasi-mapping mode, using the same index as for the ONT libraries, and including adjustments for GC content and sequence bias. Abundances were read into R using tximport (v1.8.0) ^33^. In addition, we used StringTie to assemble new transcripts (without the -e flag, provided with the reference gtf file) for comparison with the transcripts identified by FLAIR from the ONT libraries. For this analysis, we merged the HISAT2 bam files from all four Illumina samples to use as the input for StringTie. We used the default coverage cutoff of 2.5 to determine which assembled transcripts to retain in the output file.

### Public data

In addition to the ONT and Illumina data generated in-house, we processed the SIRV E0 (SRA accession number SRR6058584) and ERCC Mix1 (SRA accession number SRR6058582) ONT dRNA libraries from Garalde *et al.* ^15^. The reads were aligned to the respective transcriptomes using minimap2 with the same settings as above. The SIRV data set was also aligned to the corresponding genome using minimap2 with the settings described above, and additionally setting `—spiice-fiank=no` to accommodate the non-canonical splice sites present in this data.

## Acknowledgements

The authors would like to thank Botond Sipos, Michael Stadler and Giovanni d’Ario for constructive discussions and feedback on the manuscript. Research in the S.H. laboratory is funded by the Biotechnology and Biosciences Research Council, UK. C.S. was supported by a Pilot Project grant from the University Research Priority Program Evolution in Action of the University of Zurich. M.D.R. acknowledges support from the University Research Priority Program Evolution in Action at the University of Zurich and the Swiss National Science Foundation (310030_175841).

## Author contributions

C.S.: Conceptualization, Data curation, Formal analysis, Funding acquisition, Methodology, Software, Visualization, Writing – original draft, Writing – review & editing. Y.Y.: Formal analysis. A.B-N.: Methodology, Writing – review & editing. A.P.: Methodology. M.D.R.: Conceptualization, Data curation, Formal analysis, Funding acquisition, Methodology, Writing – review & editing. S.H.: Conceptualization, Funding acquisition, Investigation, Methodology, Writing – original draft, Writing – review & editing

## Competing interests

The authors declare that they have no competing interests.

## Data availability

The raw sequence files have been uploaded to ArrayExpress under accession numbers E-MTAB-7757 (Illumina) and E-MTAB-7778 (ONT).

## Code availability

The code used to perform the analyses in the paper is available on GitHub: https://github.com/csoneson/NativeRNAseqComplexTranscriptome.

## References

1. Keren, H., Lev-Maor, G. & Ast, G. Alternative splicing and evolution: diversification, exon definition and function. Nat. Rev. Genet. 11, 345–355 (2010).

2. Pan, Q., Shai, O., Lee, L. J., Frey, B. J. & Blencowe, B. J. Deep surveying of alternative splicing complexity in the human transcriptome by high-throughput sequencing. Nat. Genet. 40, 1413 (2008).

3. Wang, E. T. et al. Alternative isoform regulation in human tissue transcriptomes. Nature 456, 470–476 (2008).

4. Mercer, T. R. et al. Targeted RNA sequencing reveals the deep complexity of the human transcriptome. Nat. Biotechnol. 30, 99–104 (2011).

5. Vaquero-Garcia, J. et al. A new view of transcriptome complexity and regulation through the lens of local splicing variations. Elife 5, e11752 (2016).

6. Sharon, D., Tilgner, H., Grubert, F. & Snyder, M. A single-molecule long-read survey of the human transcriptome. Nat. Biotechnol. 31, 1009–1014 (2013).

7. Byrne, A. et al. Nanopore long-read RNAseq reveals widespread transcriptional variation among the surface receptors of individual B cells. Nat. Commun. 8, ncomms16027 (2017).

8. Seki, M. et al. Evaluation and application of RNA-Seq by MinION. DNA Res. (2018). doi:10.1093/dnares/dsy038

9. Oikonomopoulos, S., Wang, Y. C., Djambazian, H., Badescu, D. & Ragoussis, J. Benchmarking of the Oxford Nanopore MinION sequencing for quantitative and qualitative assessment of cDNA populations. Sci. Rep. 6, 31602 (2016).

10. Weirather, J. L. et al. Comprehensive comparison of Pacific Biosciences and Oxford Nanopore Technologies and their applications to transcriptome analysis. F1000Res. 6, 100 (2017).

11. Tilgner, H., Grubert, F., Sharon, D. & Snyder, M. P. Defining a personal, allele-specific, and single-molecule long-read transcriptome. Proc. Natl. Acad. Sci. U. S. A. 111, 9869–9874 (2014).

12. Gonzalez-Garay, M. L. Introduction to Isoform Sequencing Using Pacific Biosciences Technology (Iso-Seq). in Transcriptomics and Gene Regulation (ed. Wu, J.) 141–160 (Springer Netherlands, 2016).

13. Aird, D. et al. Analyzing and minimizing PCR amplification bias in Illumina sequencing libraries. Genome Biol. 12, R18 (2011).

14. Steijger, T. et al. Assessment of transcript reconstruction methods for RNA-seq. Nat. Methods 10, 1177 (2013).

15. Garalde, D. R. et al. Highly parallel direct RNA sequencing on an array of nanopores. Nat. Methods (2018). doi:10.1038/nmeth.4577

16. Chenchik, A. et al. RT–PCR Methods for Gene Cloning and Analysis. (1998).

17. Workman, R. E. et al. Nanopore native RNA sequencing of a human poly(A) transcriptome. bioRxiv 459529 (2018). doi:10.1101/459529

18. Carter, J.-M. & Hussain, S. Robust long-read native DNA sequencing using the ONT CsgG Nanopore system. Wellcome Open Res 2, 23 (2017).

19. Jain, M. et al. Nanopore sequencing and assembly of a human genome with ultra-long reads. Nat. Biotechnol. 36, 338–345 (2018).

20. Tyson, J. R. et al. MinION-based long-read sequencing and assembly extends the Caenorhabditis elegans reference genome. Genome Res. 28, 266–274 (2018).

21. Li, H. Minimap2: pairwise alignment for nucleotide sequences. Bioinformatics (2018). doi:10.1093/bioinformatics/bty191

22. Li, H. et al. The Sequence Alignment/Map format and SAMtools. Bioinformatics 25, 2078–2079 (2009).

23. Lawrence, M. et al. Software for computing and annotating genomic ranges. PLoS Comput. Biol. 9, e1003118 (2013).

24. Quinlan, A. R. & Hall, I. M. BEDTools: a flexible suite of utilities for comparing genomic features. Bioinformatics 26, 841–842 (2010).

25. Wang, L., Wang, S. & Li, W. RSeQC: quality control of RNA-seq experiments. Bioinformatics 28, 2184–2185 (2012).

26. Patro, R., Duggal, G., Love, M. I., Irizarry, R. A. & Kingsford, C. Salmon provides fast and bias-aware quantification of transcript expression. Nat. Methods 14, 417–419 (2017).

27. Liao, Y., Smyth, G. K. & Shi, W. The Subread aligner: fast, accurate and scalable read mapping by seed-and-vote. Nucleic Acids Res. 41, e108 (2013).

28. Liao, Y., Smyth, G. K. & Shi, W. featureCounts: an efficient general purpose program for assigning sequence reads to genomic features. Bioinformatics 30, 923–930 (2014).

29. Martin, M. Cutadapt removes adapter sequences from high-throughput sequencing reads. EMBnet.journal 17, 10–12 (2011).

30. Dobin, A. et al. STAR: ultrafast universal RNA-seq aligner. Bioinformatics 29, 15–21 (2013).

31. Pertea, M. et al. StringTie enables improved reconstruction of a transcriptome from RNA-seq reads. Nat. Biotechnol. 33, (2015).

32. Kim, D., Langmead, B. & Salzberg, S. L. HISAT: a fast spliced aligner with low memory requirements. Nat. Methods 12, 357 (2015).

33. Soneson, C., Love, M. I. & Robinson, M. D. Differential analyses for RNA-seq: transcript-level estimates improve gene-level inferences. F1000Res. 4, 1521 (2015).

