## Supplementary Figures for "A comprehensive examination of Nanopore native RNA sequencing for characterization of complex transcriptomes"

Charlotte Soneson[1,2,*,†], Yao Yao[1,2], Anna Bratus-Neuenschwander[3], Andrea Patrignani[3], Mark D. Robinson[1,2,*], Shobbir Hussain[4,*]

[1] Institute of Molecular Life Sciences, University of Zurich, 8057 Zurich, Switzerland

[2] SIB Swiss Institute of Bioinformatics, 8057 Zurich, Switzerland

[3] Functional Genomics Centre Zurich, ETHZ/University of Zurich, 8057 Zurich, Switzerland

[4] Department of Biology and Biochemistry, University of Bath, Bath BA2 7AY, United Kingdom

[†] Current affiliation: Friedrich Miescher Institute for Biomedical Research and SIB Swiss Institute of Bioinformatics, Basel, Switzerland

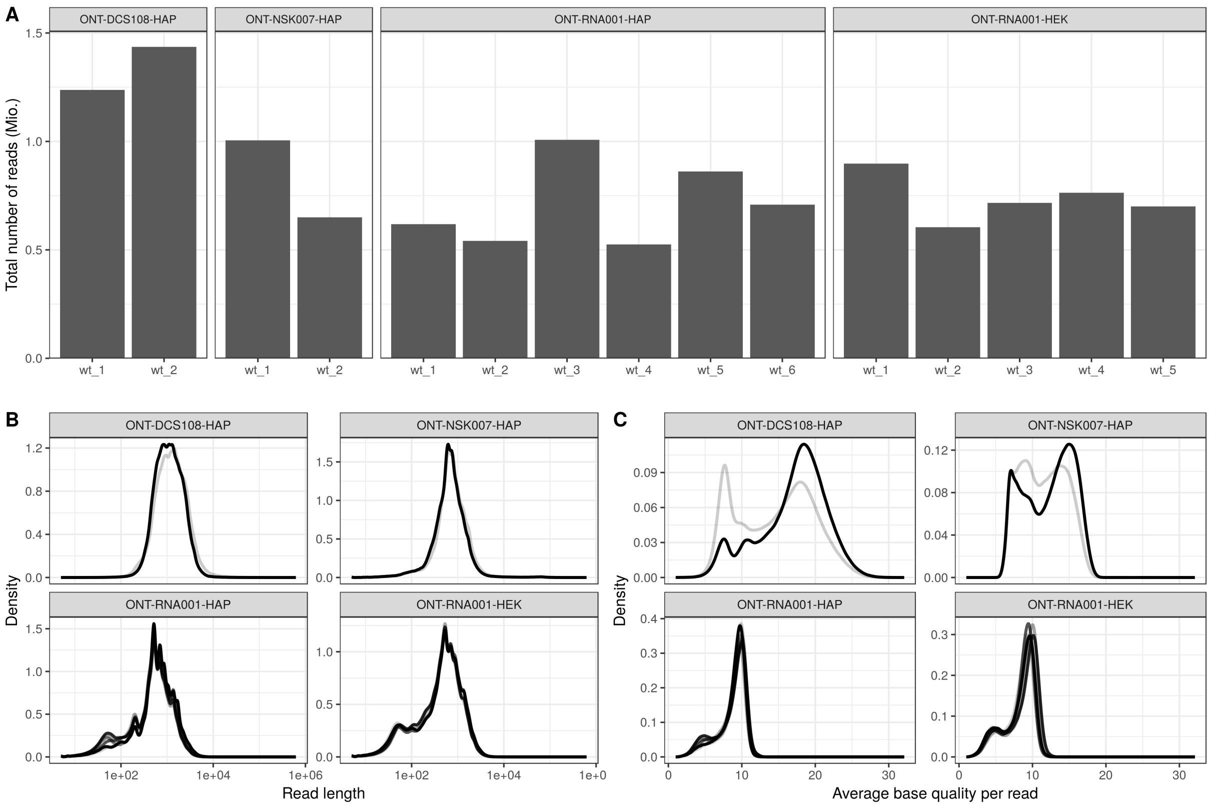


Supplementary Figure 1: Summary of the full set of reads in each ONT library, stratified by data set. A. Total number of reads. B. Read length distribution. C. Average base quality distribution. In B-C, each line corresponds to one library.

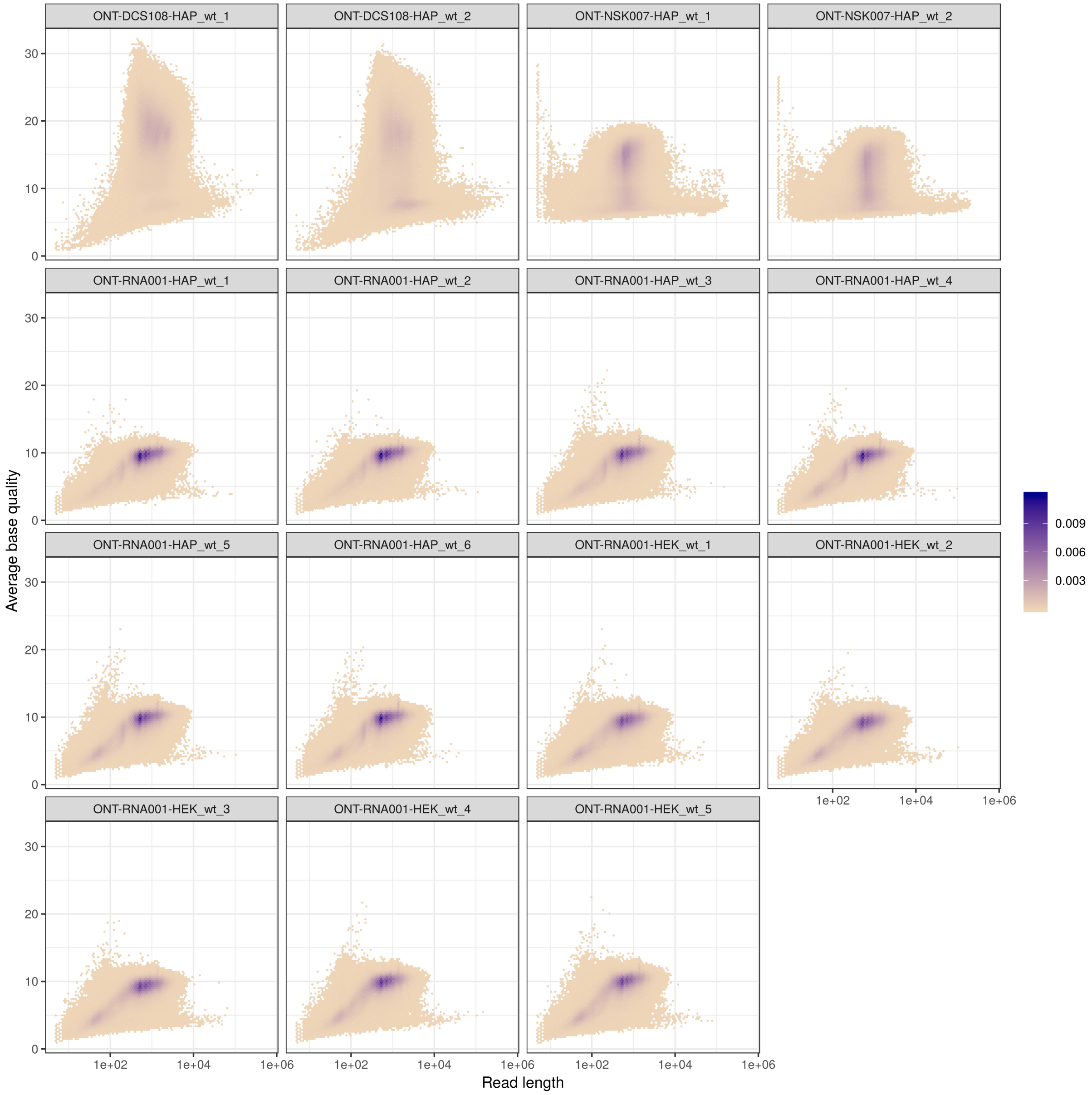


Supplementary Figure 2: Total read length ($x$) vs average base quality ($y$) for all reads in each of the ONT libraries. The colour indicates point density.

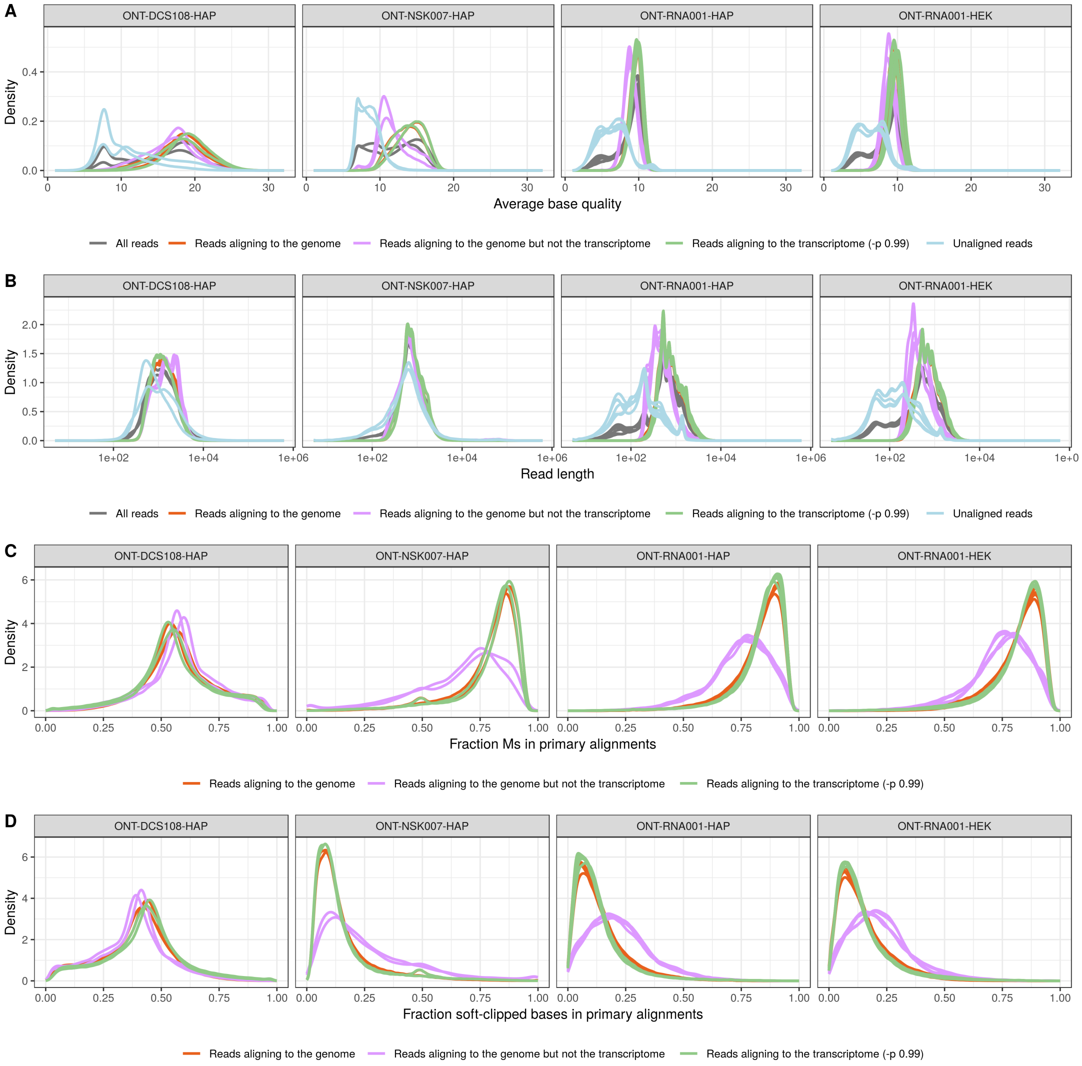


Supplementary Figure 3: Summary of read properties by alignment status. A. Average base quality distribution. Unaligned reads are predominantly found among reads with low base quality. B. Read length distribution. C. Number of "M"s in the CIGAR string of the primary alignment to the genome and transcriptome, respectively, divided by the read length. D. Number of soft-clipped ("S") bases in the CIGAR string of the primary alignment to the genome and transcriptome, respectively, divided by the read length.

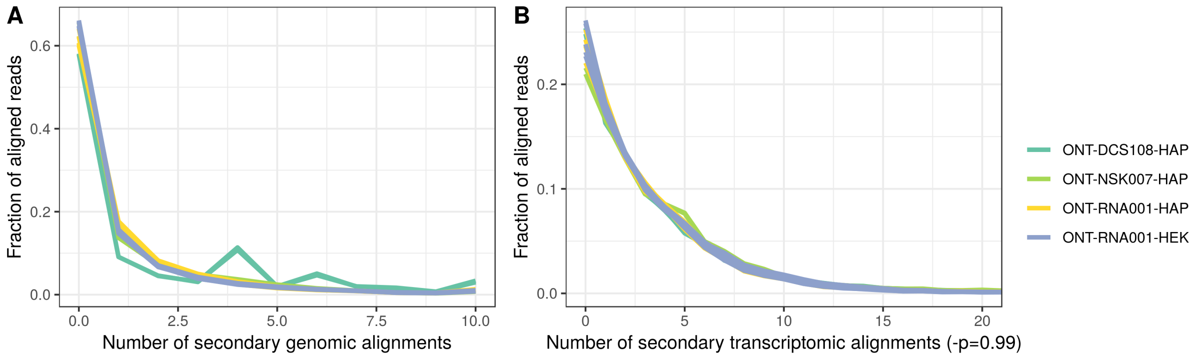


Supplementary Figure 4: Distribution of the number of secondary genome and transcriptome (with the `-p 0.99` setting) minimap2 alignments, for each library. A. For the genome alignments, up to 10 secondary alignments were allowed. B. For the transcriptome alignments, up to 100 secondary alignments were allowed, in order to allow for the high similarity among isoforms. However, very few reads had more than 20 secondary alignments, and thus the x-axis is zoomed in to the range [0-20].

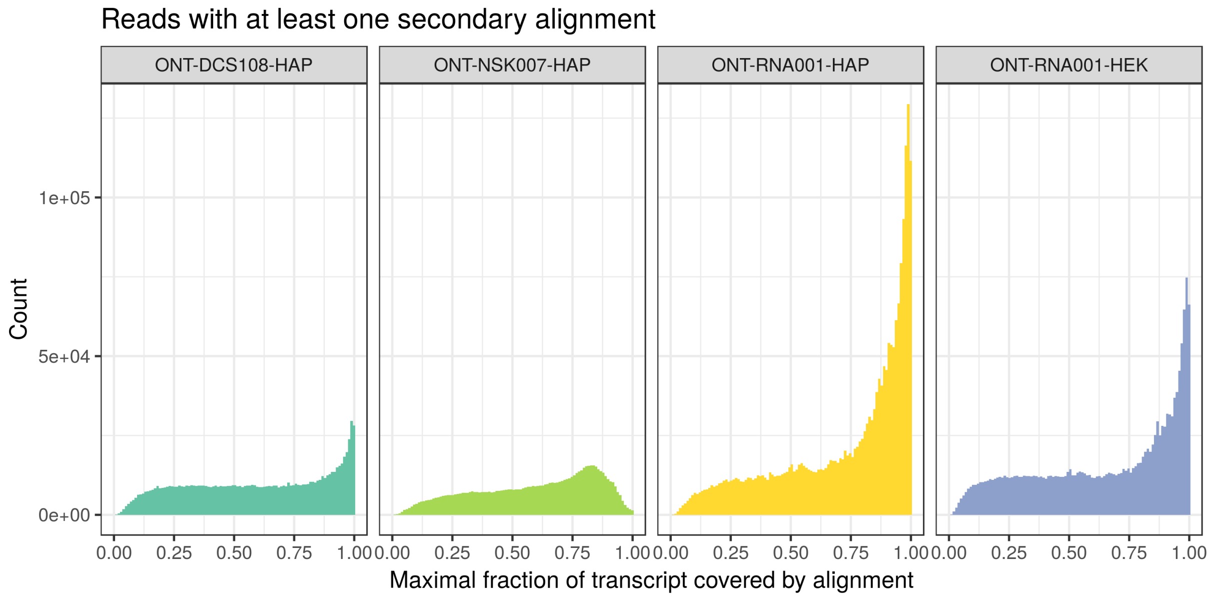


Supplementary Figure 5: The highest degree of coverage of a reference transcript by single ONT reads, for each read with at least one secondary alignment.

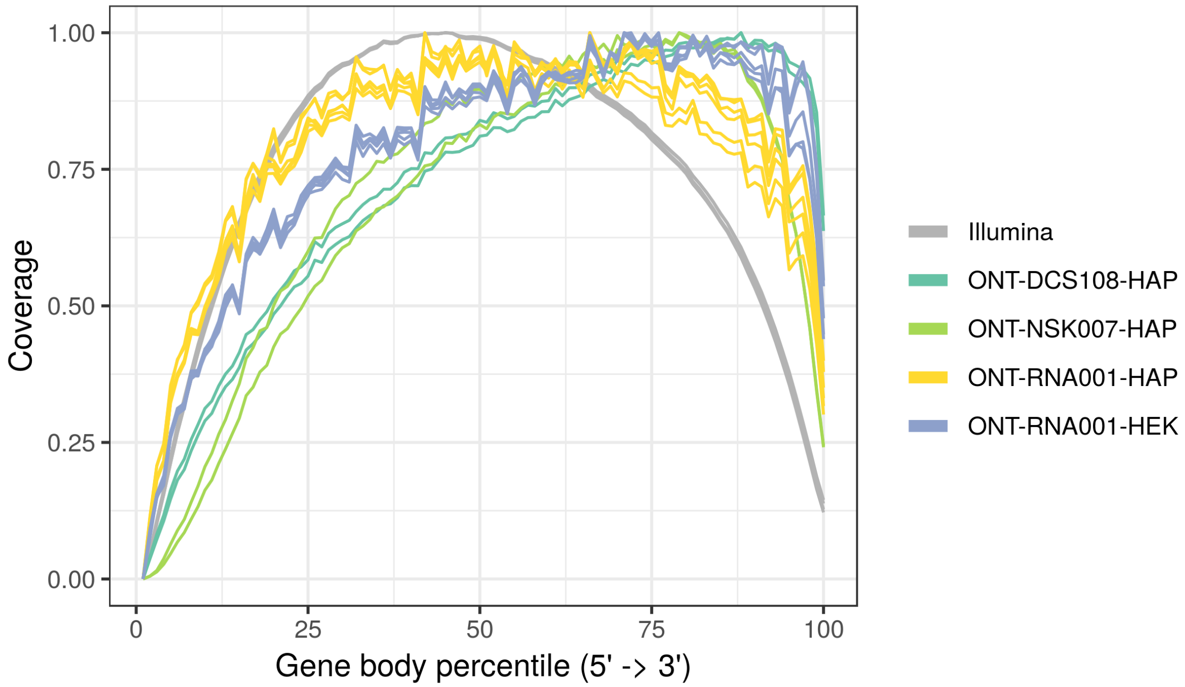


Supplementary Figure 6: Gene body coverage, estimated by RSeQC, in the ONT and Illumina libraries.

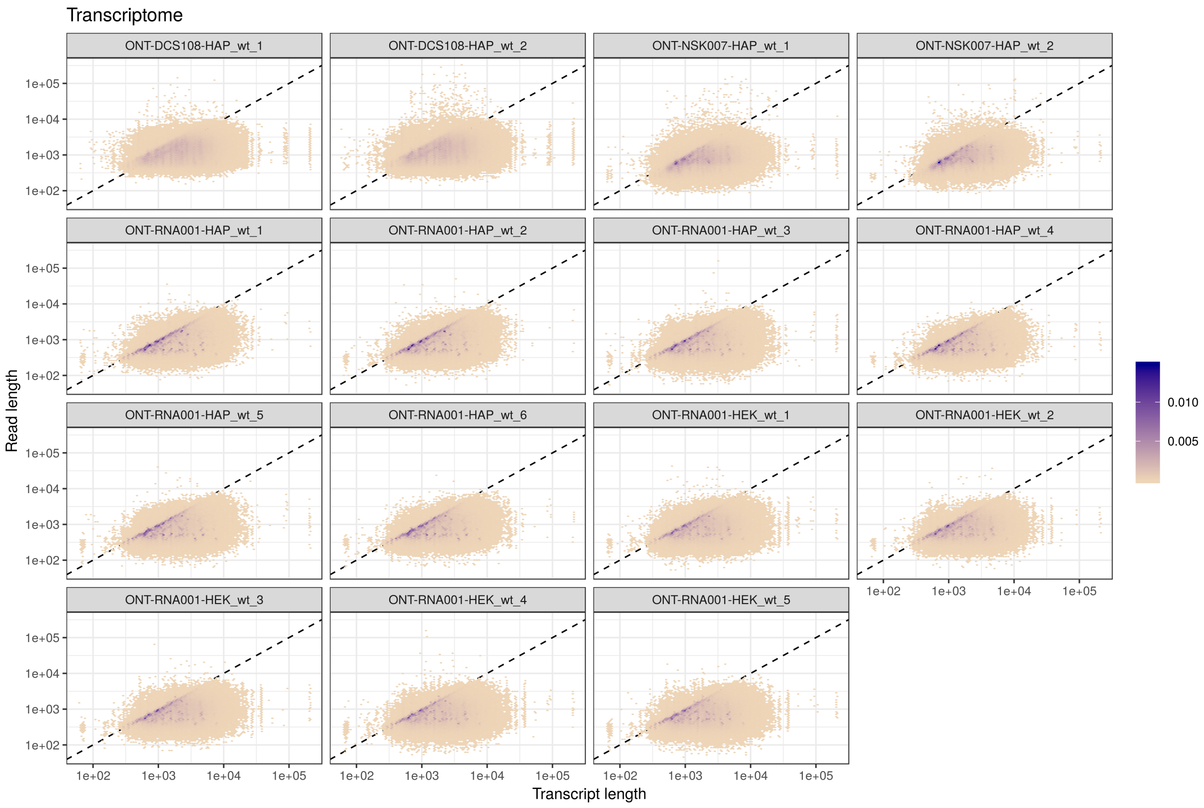


Supplementary Figure 7: Annotated length of the target transcript ($x$) vs total read length ($y$) for all primary transcriptome alignments in each of the ONT libraries. Reads aligning to the shortest transcripts are often longer than the transcript, and are thus soft clipped in the alignment. The colour indicates point density.

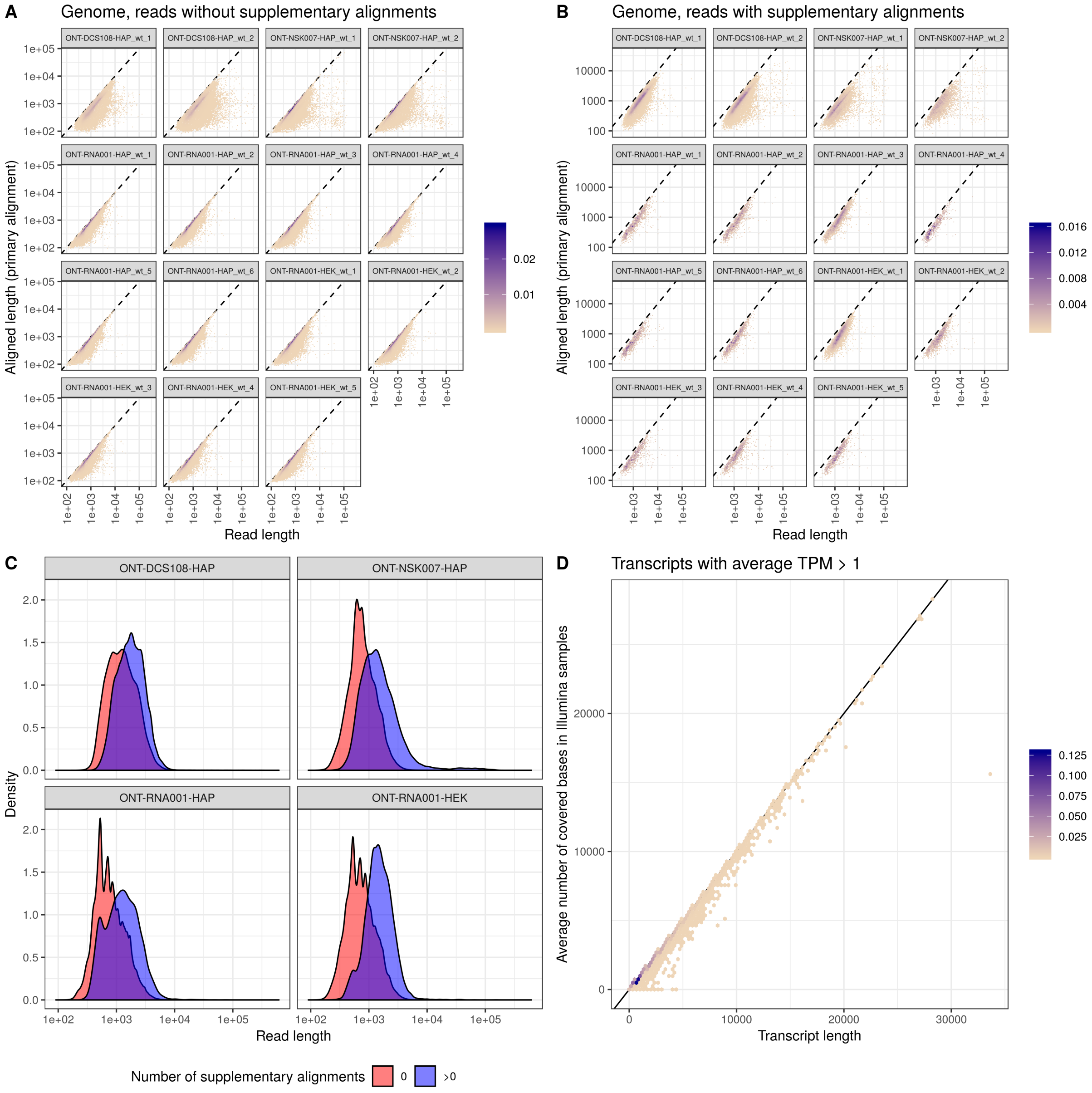


Supplementary Figure 8: A-B. Total read length ($x$) vs length of the primary genome alignment ($y$) for reads without (A) and with (B) any reported supplementary alignment. The colour indicates point density. C. Read length distribution for reads without (red) and with (blue) any reported supplementary alignment, in the four ONT data sets. D. The average number of nucleotides annotated to the transcript that are covered by reads in the Illumina samples *vs* the transcript length, for transcripts with an average estimated TPM exceeding 1 across the Illumina samples.

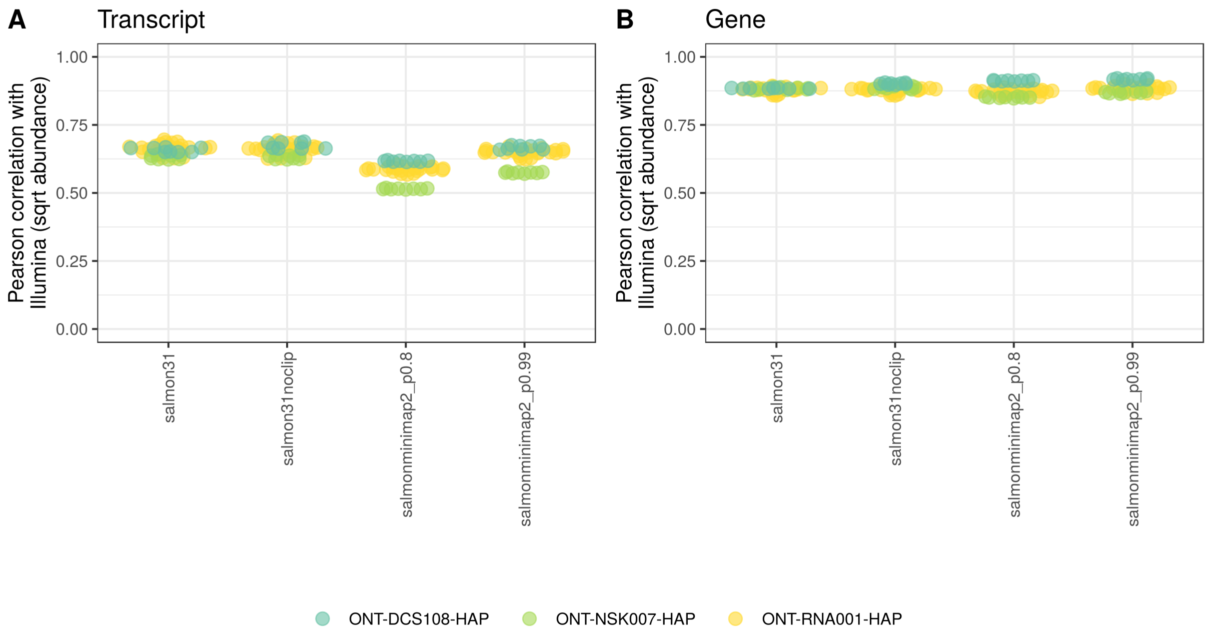


Supplementary Figure 9: Correlation between estimated abundances in the Illumina samples and those obtained from the ONT data, running Salmon in different configurations. ONT abundances are estimated counts, and Illumina abundances are estimated TPMs from Salmon. Each point corresponds to a pair of one Illumina sample and one ONT sample, and all such pairwise combinations were considered.

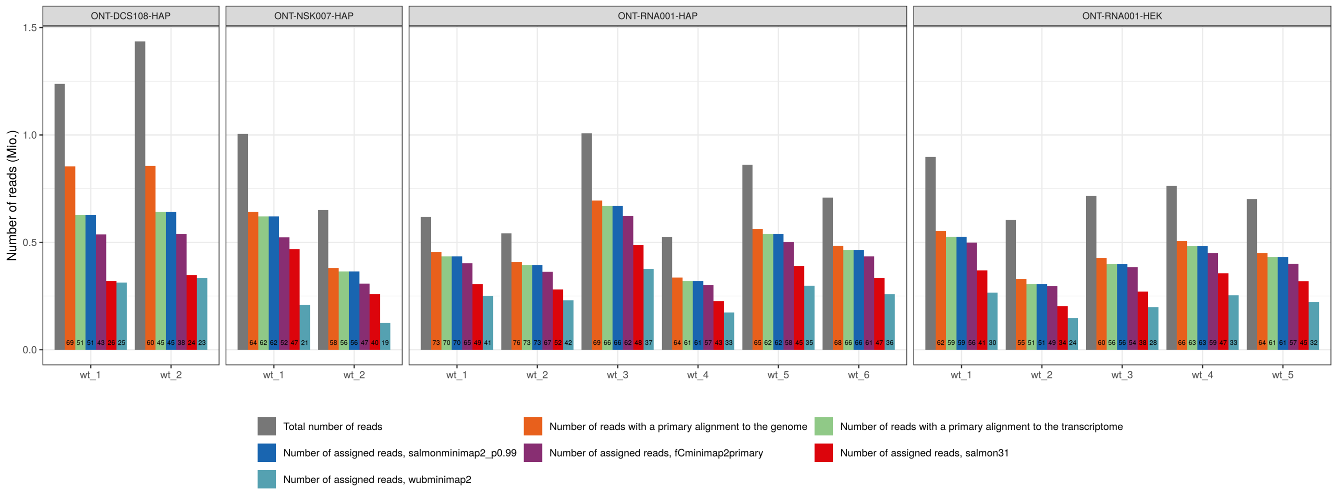


Supplementary Figure 10: Number of reads assigned to features (genes or transcripts) by each of the abundance estimation methods, for each ONT library. The number written in each bar indicates the percentage of the total number of reads that are assigned to features by the corresponding method.

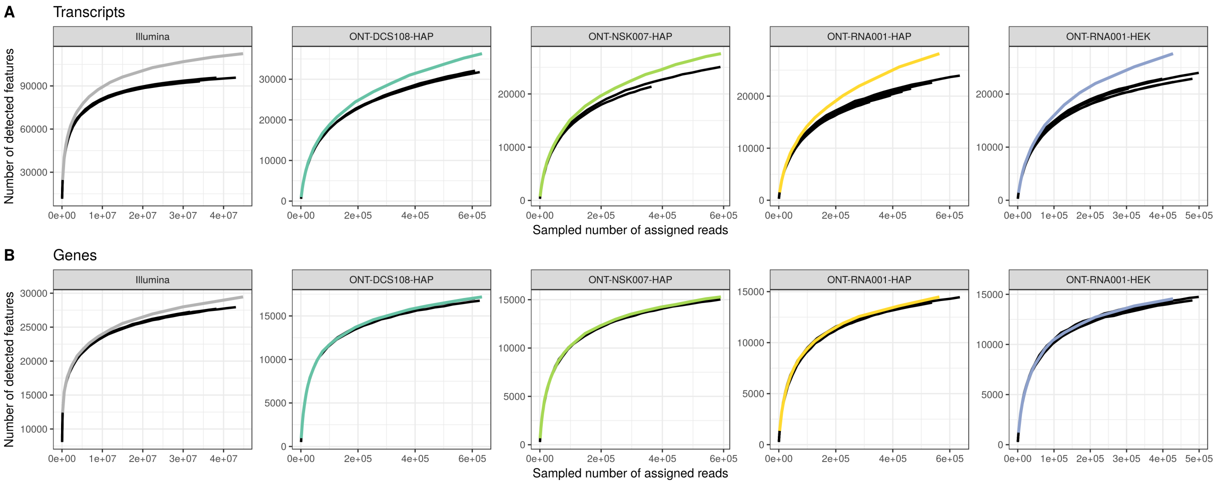


Supplementary Figure 11: Saturation curves for transcript (A) and gene (B) detection. Black curves represent individual libraries, and colored curves are obtained by pooling the reads across all samples in the data set before subsampling. These curves are truncated to the same range as the individual sample curves to facilitate comparison. A feature is considered detected if it has an estimated salmonminimap2 count (ONT libraries) or Salmon count (Illumina libraries) $\geq 1$.

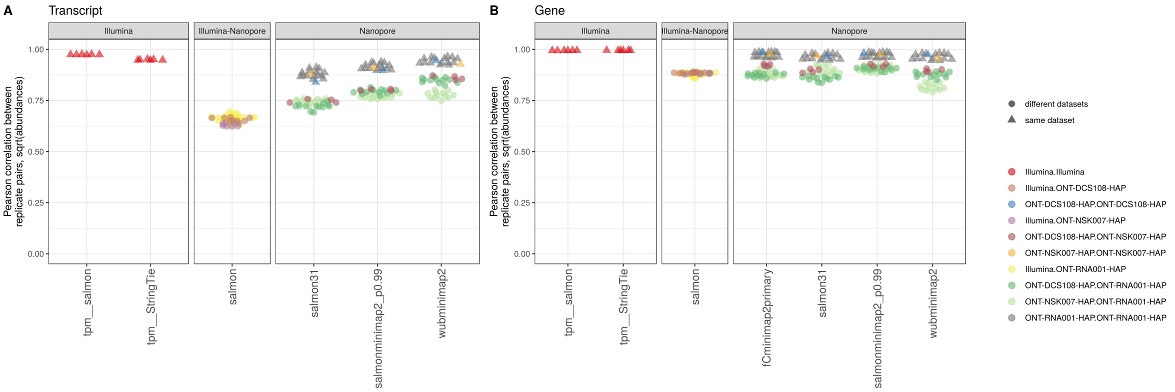


Supplementary Figure 12: Pearson correlations between transcript (A) or gene (B) abundances for each pair of samples, within and between data sets, for each abundance estimation method. Triangles correspond to pairs of samples within the same data set, and circles to pairs from different data sets. Correlations were calculated between square root-transformed estimated counts from the respective ONT methods, and for square root-transformed estimated TPMs for the Illumina libraries.

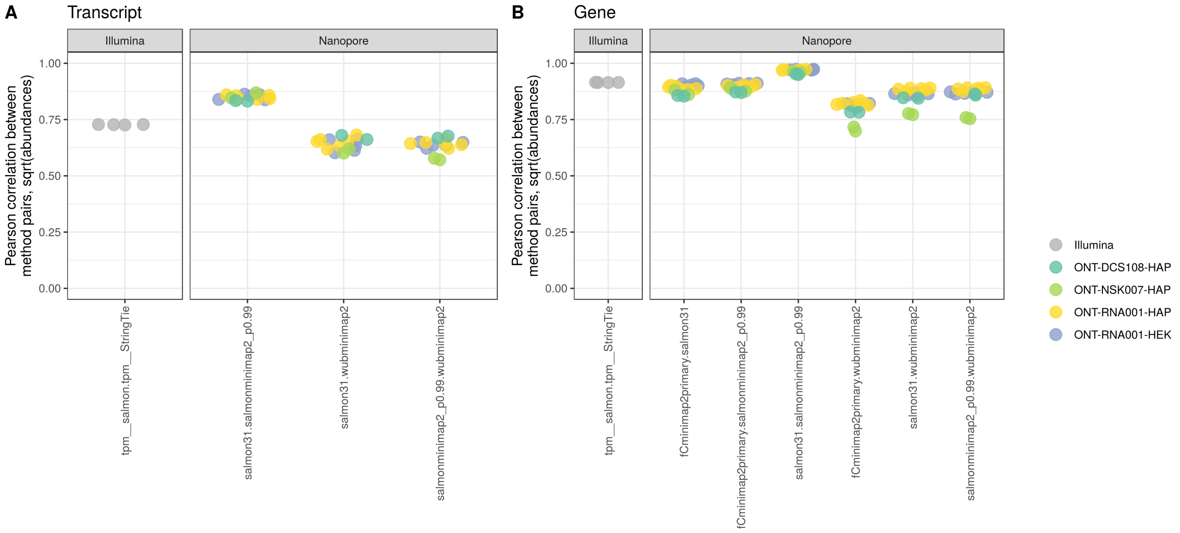


Supplementary Figure 13: Pearson correlations between transcript (A) or gene (B) abundances for each pair of abundance estimation methods, for each library. Correlations were calculated between square root-transformed estimated counts from the respective ONT methods, and for square root-transformed estimated TPMs for the Illumina libraries.

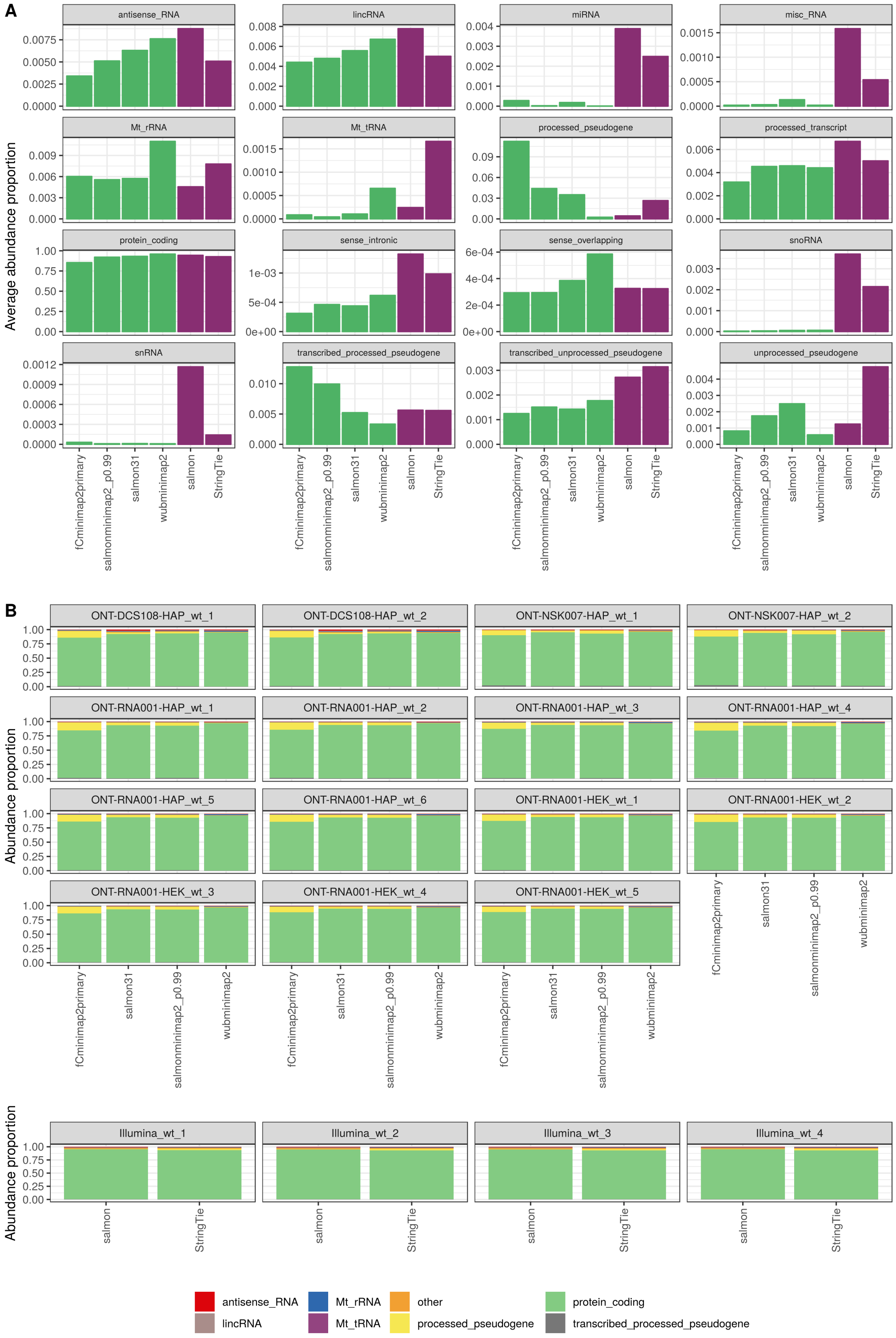


Supplementary Figure 14: A. Relative abundance (proportion of total abundance) assigned to genes of different biotypes by the different quantification methods, averaged across all libraries. Green bars represent abundances (counts) for ONT data, while purple bars represents abundances (TPMs) for the Illumina libraries. B. Relative abundance (proportion of total count) assigned to genes of different biotypes by the different quantification methods, for all libraries. Only the most abundant biotypes are represented separately, all others are collapsed into the "other" category.

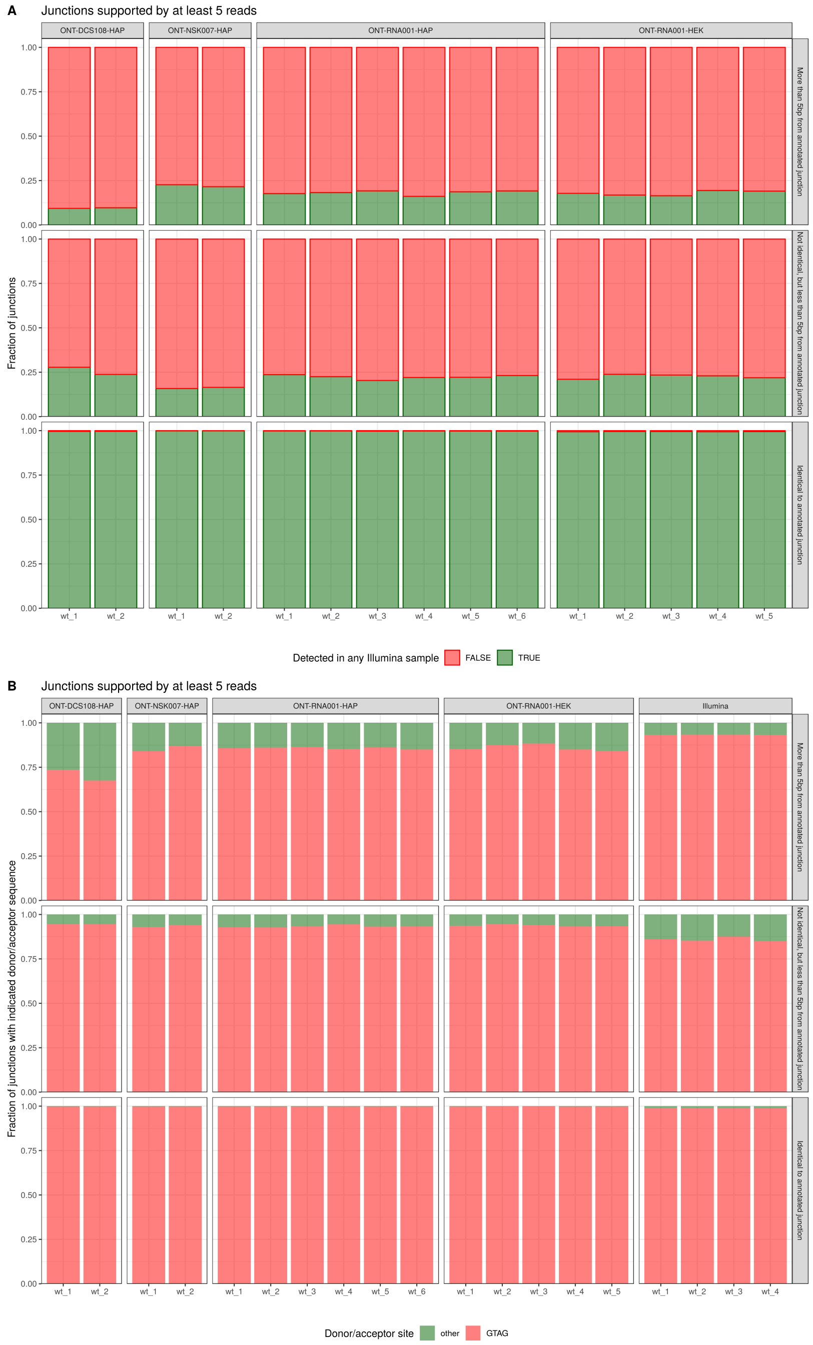


Supplementary Figure 15: A. Fraction of the junctions covered by at least 5 ONT reads that are also detected in at least one of the Illumina samples (supported by at least 1 read), for each category. B. Fraction of the junctions that contain a canonical (GT-AG) or non-canonical splicing motif, respectively, for each category.

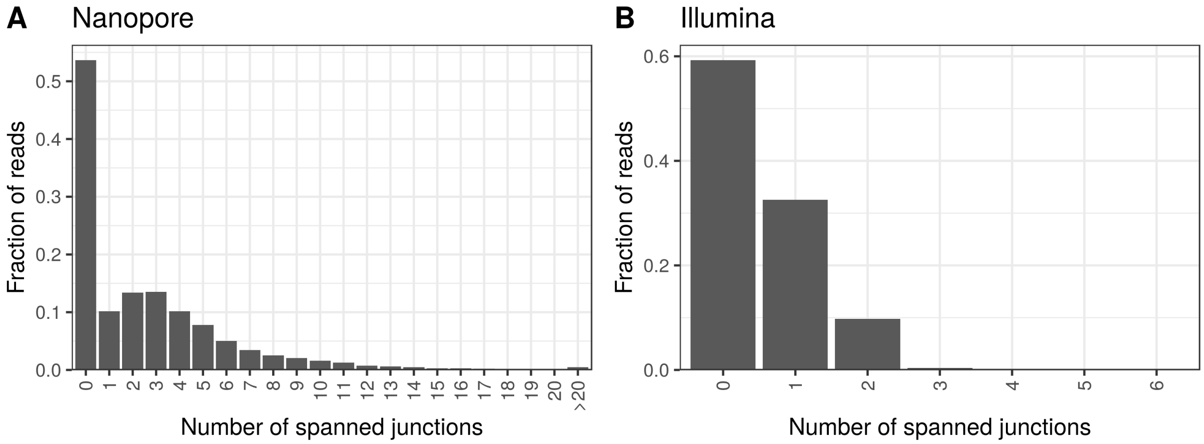


Supplementary Figure 16: Distribution of the number of junctions spanned by individual reads in the ONT (A) and Illumina (B) libraries. For each Illumina library, the number of spanned junctions were counted for a subset of 5 million reads, and only reads being the first in a properly mapped pair were contributing to the summary shown in this plot.

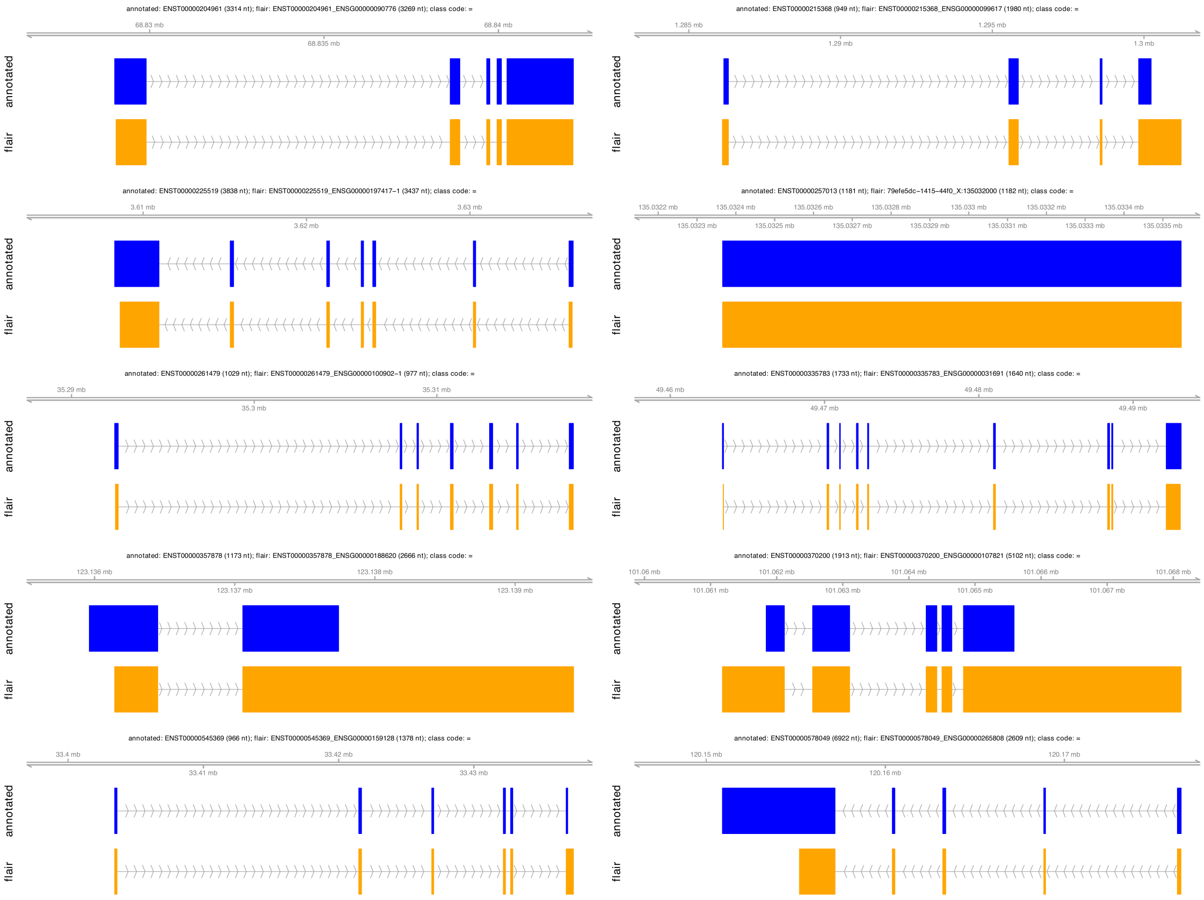


Supplementary Figure 17: Examples of transcripts identified by FLAIR in the *ONT-RNA001-HAP* data set, with a complete, exact match to the intron chain of an annotated transcript (class code '=' from gffcompare).
